# A Novel Regioselective Approach to Cyclize Phage-Displayed Peptides in Combination with Epitope-Directed Selection to Identify a Potent Neutralizing Macrocyclic Peptide for SARS-CoV-2

**DOI:** 10.1101/2022.07.06.498864

**Authors:** J. Trae Hampton, Tyler J. Lalonde, Jeffery M. Tharp, Yadagiri Kurra, Yugendar R. Alugubelli, Christopher M. Roundy, Gabriel L. Hamer, Shiqing Xu, Wenshe Ray Liu

**Affiliations:** Texas A&M Drug Discovery Laboratory, Department of Chemistry, Texas A&M University, College Station, TX 77843, USA; Department of Entomology, Texas A&M University, College Station, TX 77843, USA; Institute of Biosciences and Technology and Department of Translational Medical Sciences, College of Medicine, Texas A&M University, Houston, TX 77030, USA; Department of Biochemistry and Biophysics, Texas A&M University, College Station, TX 77843, USA; Department of Molecular and Cellular Medicine, College of Medicine, Texas A&M University, College Station, TX 77843, USA

**Keywords:** Phage display, macrocyclic peptides, Asymmetric cyclization, SARS-CoV-2, antiviral

## Abstract

Using the regioselective cyanobenzothiazole condensation reaction with the *N*-terminal cysteine and the chloroacetamide reaction with an internal cysteine, a phage-displayed macrocyclic 12-mer peptide library was constructed and subsequently validated. Using this library in combination with iterative selections against two epitopes from the receptor binding domain (RBD) of the SARS-CoV-2 Spike protein, macrocyclic peptides that strongly inhibit the interaction between the Spike RBD and ACE2, the human host receptor of SARS-CoV-2, were identified. The two epitopes were used instead of the Spike RBD to avoid selection of nonproductive macrocyclic peptides that bind RBD but do not directly inhibit its interactions with ACE2. Antiviral tests against SARS-CoV-2 showed that one macrocyclic peptide is highly potent against viral reproduction in Vero E6 cells with an EC_50_ value of 3.1 μM. The AlphaLISA-detected IC_50_ value for this macrocyclic peptide was 0.3 μM. The current study demonstrates that two kinetically-controlled reactions toward *N*-terminal and internal cysteines, respectively, are highly effective in the construction of phage-displayed macrocyclic peptides, and the selection based on the SARS-CoV-2 Spike epitopes is a promising methodology in the identification of peptidyl antivirals.

## INTRODUCTION

With the potential to combine the high specificity of large antibodies and the stability of small molecules, macrocyclic peptides occupy a promising area of drug discovery.^1^ Because of this, many different techniques have been developed to screen macrocyclic peptides, ranging from phage display to combinatorial chemistry.^2–6^ To date, many methods have been developed to chemically cyclize peptides displayed on the phage surface.^3,4,7^ The classical approach took advantage of the disulfide formation between two cysteines that flank the peptide library on both sides (Figure 1A).^8^ Although this has been used to identify macrocyclic peptide ligands, their translation into therapeutics is problematic due to the instability of disulfide bonds under the reducing environment in cells.^9^ To counter this, cysteine-reactive bis-electrophiles have been developed to cyclize phage-displayed peptides containing two flanking cysteines via thioether bond formation (Figure 1B). An extension of this idea has also led to the development of phage-displayed bicyclic peptides by chemical cyclization of three cysteine in a phage-displayed peptide.^3^ These linkers typically contain activated halogens that will react with any cysteine on the phage.^4^ However, this low specificity often results in phage toxicity at high concentrations, thus decreasing the efficiency of selections.^3^ To avoid this, our group and others have developed genetically encoded cyclic libraries that take advantage of proximity-driven reactions between a genetically encoded noncanonical amino acid and a cysteine, yet these also result in low phage yields compared to libraries only containing canonical amino acids.^2,10^ Due to the aforementioned limitations of current cyclization approaches of phage-displayed peptides, a technique that can efficiently cyclize phage-displayed peptides but limit modifications of non-targeted phage residues is still desired. For this purpose, we aimed to develop a highly specific bis-electrophile that preferentially reacts with an *N*-terminal cysteine by taking advantage of the reaction between cyanobenzothiazole (CBT) and 1,2-aminothiols, which is the last step in firefly luciferin biosynthesis.^11^ This condensation proceeds rapidly under physiological conditions with a second order rate constant of 9.19 M^-1^ s^-1^,^12^ and has been used for a variety of biological techniques, ranging from protein labelling to nanoparticle self-assembly.^12–14^ Thus, by fusing the cyanobenzothiazole core to a chloroacetamide moiety to generate *N*-chloroacetyl 2-chloro-*N*-(2-cyanobenzo[*d*]thiazol-6-yl)acetamide (CAmCBT), we envisioned it would result in a nontoxic cyclic peptide linker for phage display that preferentially reacts with an *N*-terminal cysteine, followed by a proximity-driven SN_2_ reaction with an internal cysteine (Figure 1B). In this paper, we report our progress in the successful construction of a phage-displayed peptide library using this proposed approach and its application in the identification of a macrocyclic peptide antiviral for SARS-CoV-2.

**Figure 1:**
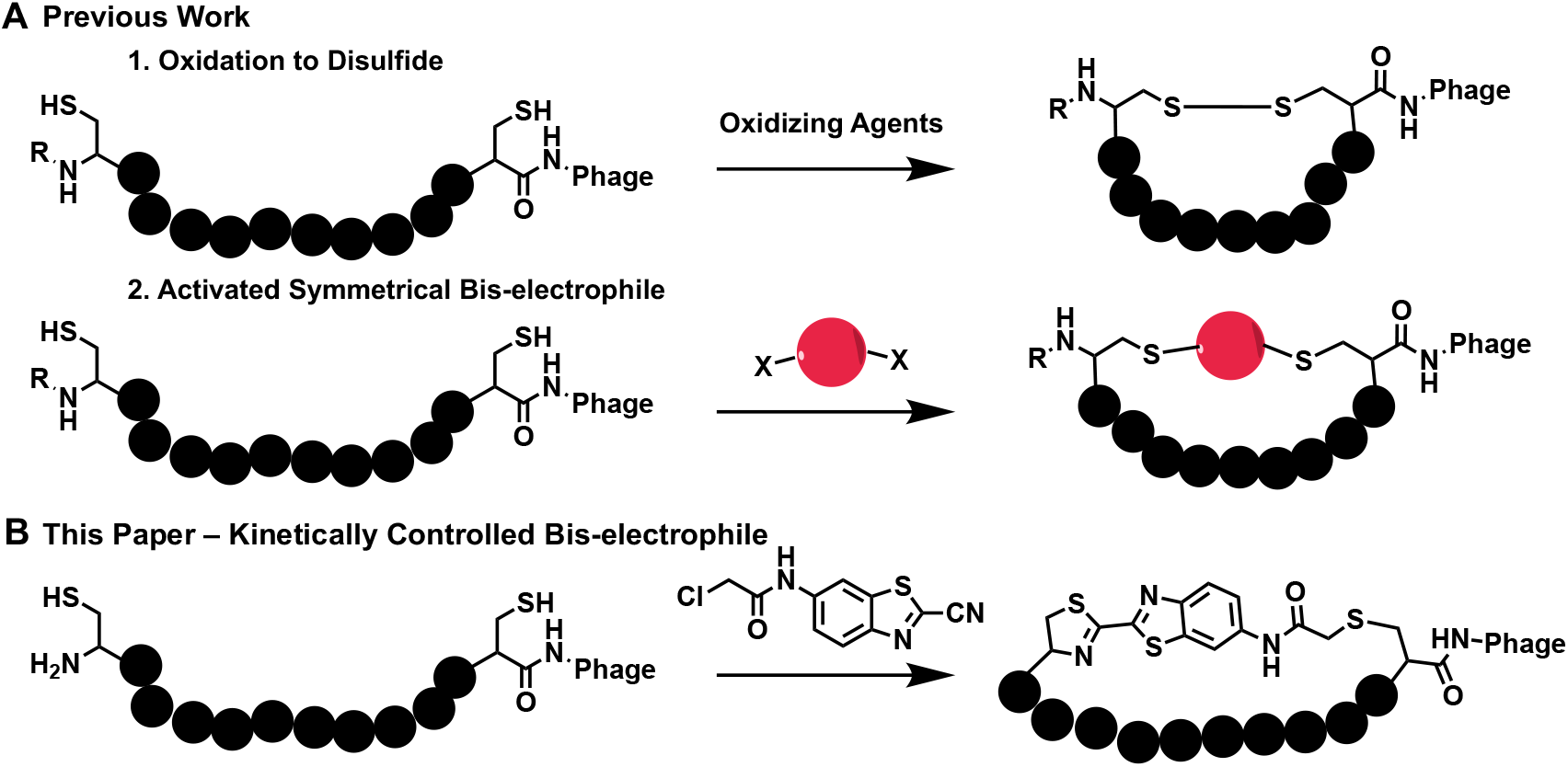
Current cyclic approaches for phage-displayed peptides and our proposed approach. **A)** Previous methods for the cyclization of two cysteines in phage-displayed peptides. R = H or peptide chain. **B)** The proposed novel cyclic linker CAmCBT that provides an asymmetric cyclization scaffold for phage display by selectively reacting with the *N*-terminal cysteine and then a proximity-driven reaction with an internal cysteine.

SARS-CoV-2 is the viral pathogen for COVID-19 that has been a global pandemic with more than half a billion confirmed cases and over 6 million deaths. While there are efficacious vaccines that have been developed that show promise in multiple variants of concern, there is still a need for treatments that neutralize the virus for both breakthrough infections and places where vaccine distribution has been hindered. Antivirals that include small molecules, peptides and antibodies are also essential for treating patients that show severe symptoms. There are several demonstrated routes for the development of SARS-CoV-2 antivirals by targeting different essential proteins. The SARS-CoV-2 genome encodes for 12 open reading frames (ORFs), with four of these coding structural proteins for virion formation: the Spike glycoprotein, a membrane protein, a nucleoprotein, and an envelope protein.^15^ The cellular entry of SARS-CoV-2 is mediated through interactions between the Spike protein and angiotensin-converting enzyme 2 (ACE2) in the human host cell.^16^ The Spike protein engages ACE2 using its receptor binding domain (RBD). Blocking the interactions between RBD and ACE2 is a validated route for the development of SARS-CoV-2 antivirals, as several RBD-targeting antibodies and small proteins have been approved for the treatment of COVID-19.^17–25^ Compared to antibodies, small molecules and peptides have advantages in their easy manufacturing, storage and administration.^1,26^ Technically, both small molecule and peptide inhibitors of RBD-ACE2 interactions can be identified by searching ligands that bind to RBD. Several precedent works that used RBD directly for the identification of peptide ligands have been reported; however, many of the identified ligands bound the RBD at locations distal to the RBD-ACE2 binding interface, and were therefore not productive at inhibiting the RBD-ACE2 interaction.^27,28^ To avoid the identification of nonproductive peptide ligands, we reasoned that using small RBD epitopes that directly interact with ACE2 for the selection of phage-displayed macrocyclic peptides will increase the chances of identifying productive macrocyclic peptide ligands that directly block interactions between RBD and ACE2. In this work, we report the successful implementation of this approach for the identification of a potent peptidyl SARS-CoV-2 antiviral and its validation.

## RESULTS & DISCUSSION

### Development of a cyanobenzothiazole-based macrocyclic peptide linker for phage display

A variety of organic linkers have been developed to produce macrocyclic peptide libraries on phage, yet there are several limitations that hinder the diversity of the libraries they produce. Highly-reactive groups, such as benzyl bromides and bromoacetamides have been used to ensure reaction rates high enough to be practical for cyclizing phage-displayed peptides.^4,29^ Also, to avoid any complications with formation of different stereoisomers during the cyclization, traditional linkers have needed to have two-fold or three-fold symmetry. Recently, the Mayer group reported an asymmetric cyclization strategy by taking advantage of alkylation of an internal cysteine, followed by reductive amination of the N-terminus.^30^ While this approach has been innovative in providing asymmetric scaffolds, it is hindered by long reaction times (>24 h). Therefore, we looked to create an alternative asymmetric linker that has quick reactivity and regioselectivity.

2-amino-cyanobenzothiazole (ACBT) has been previously shown to quickly undergo condensation with an *N*-terminal cysteine, and other groups have modified the amine to a variety of moieties that allow for multiple applications in biology, such as tagging proteins with fluorophores or biotin.^12^ Thus, we aimed to create a cyclic linker for phage display by converting the amine of ACBT to a chloroacetamide that could react with an internal cysteine following the condensation reaction. Chloroacetamides react slowly with cysteines at neutral pH, with second-order rate constants less than 0.02 M^-1^s^-1^.^31^ This reaction rate is 450-fold slower than the condensation reaction between a cyanobenzothiazole and an *N*-terminal cysteine, which has a second-order rate constant of 9.19 M^-1^s^-1^. We envisioned that the cyclization of phage-displayed peptides flanked by an *N*-terminal cysteine and an internal cysteine using CAmCBT would afford a quick CBT condensation reaction with the *N*-terminal cysteine, and then a proximity-driven reaction between the internal cysteine with the chloroacetamide of CAmCBT. The rapid CBT condensation with the *N*-terminal cysteine will create spatial proximity between the internal cysteine and the chloroacetamide moiety of CAmCBT to facilitate quick peptide cyclization. Since the second step S_N_2 reaction is accelerated by the first condensation reaction by bringing two reactive groups into proximity, in theory only a low concentration of CAmCBT will be required for the cyclization process, ensuring that nonselective reactions with other phage cysteines or lysines can be avoided. As a model protein to test the hypothesis, a superfolder green fluorescent protein (sfGFP) with an *N*-terminal CA_5_C peptide (CA_5_C-sfGFP) was expressed. A TEV protease cutting site was introduced immediately before the *N*-terminal cysteine for the catalytic cleavage by TEV protease. A *C-*terminal 6×His tag was introduced as well for affinity purification using Ni-NTA resin. After expression in *E. coli* and purification using Ni-NTA resin, the purified protein was incubated with TEV protease to afford an exposed *N-*terminal cysteine. This was subsequently used to test the reactivity with CAmCBT. After reduction with 1 mM TCEP for 30 minutes to reduce a potential disulfide bond between the two cysteines in CA_5_C, CA_5_C-sfGFP (100 nM in PBS, pH 7.4) was reacted with 100 μM of CAmCBT for 3 hours at room temperature. At this concentration, the chloroacetamide moiety is expected to have a half-life around 100 h to label a cysteine side chain thiolate. So, with a 3-hour incubation, very minimal nonspecific labeling of cysteines should be observed. The CBT moiety was expected to react selectively with the *N*-terminal cysteine. At 100 μM, the calculated half-life for the *N*-terminal cysteine labeling is 12.5 min. Therefore, a 3-hour incubation was expected to lead to near quantitative conversion. Excitingly, electrospray ionization mass spectrometry (ESI-MS) analysis of the reacted protein indeed indicated nearly complete cyclization, with little to no uncyclized protein detected (Figure 2A). The deconvoluted ESI-MS-detected molecular weight of the reacted protein was 28,454 Da and matched well with the calculated theoretical molecular weight (28,455 Da). The nonselective chloroacetamide reaction of CAmCBT with a cysteine thiolate will lead to a protein product with molecular weight of 28,472 Da, which was not observed in the ESI-MS spectrum. Furthermore, the reacted protein had a longer resolution time than the original protein in C4 column liquid chromatography (LC) indicating it had a higher hydrophobicity due to its cyclization with CAmCBT (Figure S1).

**Figure 2:**
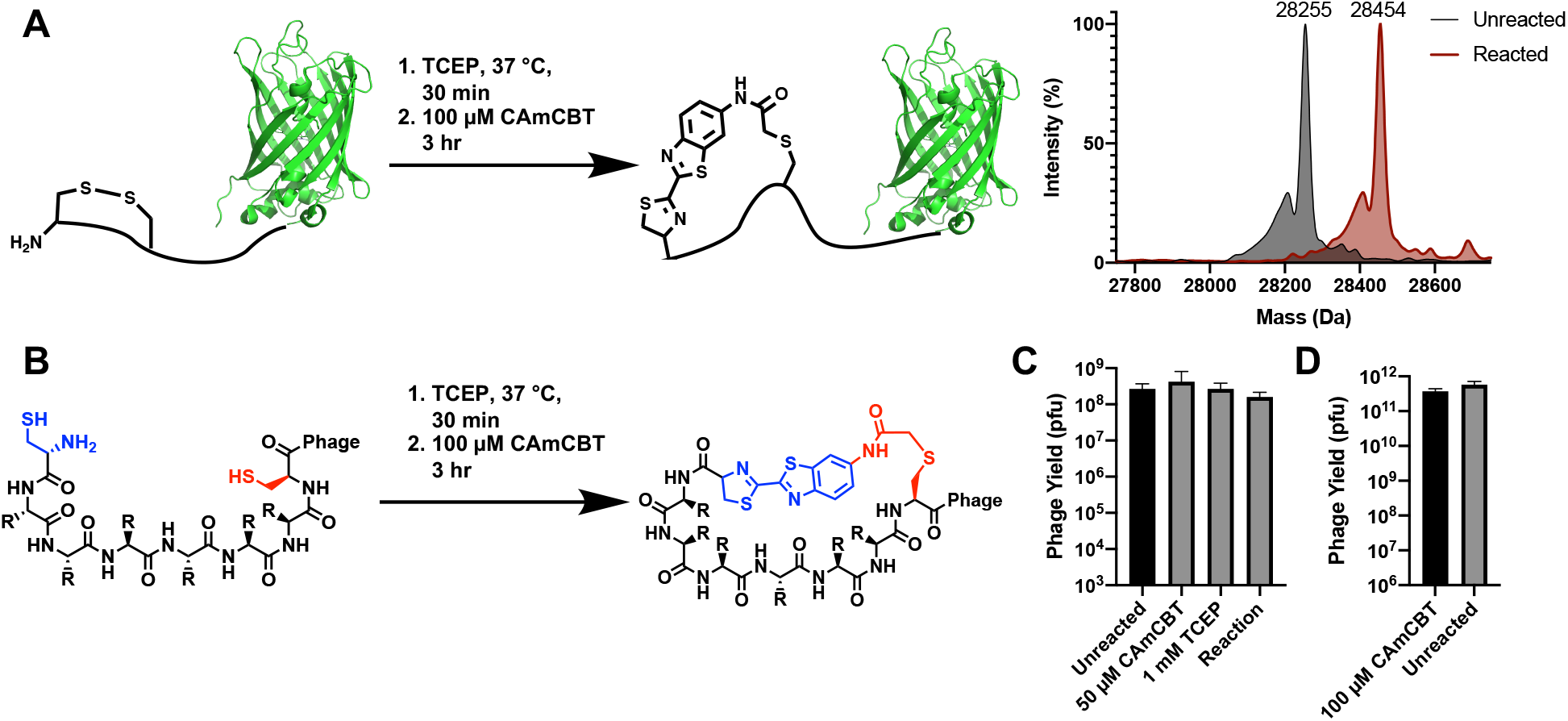
Validation of the CAmCBT cyclization linker for a peptide flanked by an *N*-terminal cysteine and an internal cysteine. **A)** Superfolder GFP (sfGFP) with an *N*-terminal CA_5_C peptide was used as a model protein to react with CAmCBT. Deconvoluted ESI-MS spectra indicated nearly complete conversion to the desired product (Expected molecular weight: 28,455 Da) after reacting for 3 h at room temperature. **B)** Treatment of phage libraries using the kinetically-controlled bis-electrophile CAmCBT. **C)** Treatment of phages containing a 5mer peptide library resulted in little to no toxicity. Phages were treated with both CAmCBT and TCEP separately before the combination of the two (Reaction) was tested. **D)** Phages containing a 12mer peptide library were reacted with 1 mM TCEP and 100 μM of CAmCBT and the reaction showed no significant phage toxicity.

Given the promising protein cyclization experiments, we proceeded to test the effectiveness and toxicity of the reaction conditions on phage. There are two endogenous cysteines in pIII, along with three additional cysteines in other phage coat proteins that could react with cysteine-reactive compounds. None of them are *N*-terminal cysteines. To ensure that the CBT condensation occurred selectively on phages containing *N*-terminal cysteines, a fluorescein-labelled CBT (FITC-CBT) was reacted with wild-type phages (M13K07) and phages that contained a 5-mer peptide library flanked by an *N*-terminal and an internal cysteine on the *N*-terminus of pIII. Only phages that contained the library were fluorescent after the reaction with FITC-CBT, thus indicating the specificity of the CBT condensation reaction for the *N*-terminal cysteine on phage (Figure S2). To confirm the reaction did not lead to any reduction in phage viability, phage infectivity was quantified before and after reaction with TCEP and CAmCBT. Incubation of the phages with 1 mM TCEP, followed by a 3 h reaction with either 50 or 100 μM of CAmCBT in PBS (pH 7.4) showed no significant reduction in phage viability, further supporting the specificity of the linker for reacting with only the *N*-terminal cysteine (Figure 2C, 2D). Phages were reacted separately with 1 mM TCEP for 30 min and tested for viability, showing no significant reduction of viability as well (Figure 2C).

### Selection of cyclic peptides that bind to SARS-CoV-2 Spike epitopes

After validation of CAmCBT for phage display, we then looked to use it to identify macrocyclic peptides that bind to the SARS-CoV-2 Spike protein and disrupt its interactions with ACE2. While there have been several peptides designed that have shown binding to the Spike RBD, there have been issues with this binding translating to inhibition of interactions with ACE2.^27,28,32^ Displayed peptides have been selected against the Spike RBD to identify peptides that block interactions between RBD and ACE2; however, however, these peptides targeted cavities of RBD that are not at the binding interface with ACE2.^28^ Therefore, rather than screening against the Spike RBD directly and potentially selecting for nonproductive binders, we used previous structural studies to design two peptide epitopes, P28 and P29, that correspond to two critical loops (residues I472-F490 and Q498-Q506, respectively) involved in binding to ACE2 (Figure 3A).^33^ These peptides were cyclized through disulfide bonds to mimic the loops in the Spike RBD, and were biotinylated to allow for immobilization for phage display screening. A 12-mer phage library that consisted of an *N*-terminal and an internal cysteine flanking 12 randomized amino acids was cyclized using the CAmCBT linker and then used for biopanning against the P28 and P29 peptides. The enrichment of phages throughout the selection was monitored through phage titers at each round. To ensure the enrichment was a result of macrocyclic phages, control selections were also performed that included the same phages without the addition of CAmCBT. After the first round of selection, more stringency was provided by increasing the number of washes. After three rounds of selection, titers indicated enrichment of phages that bound when cyclized (Figure S3). In general, macrocyclic phages were enriched 10 times better than phages without cyclization. Phage libraries from the first and third rounds of selection were isolated, and llumina paired-end sequencing was performed in a similar manner to previous work in our lab.^34^ For analysis, samples were filtered based on paired-end processing and enrichment between rounds 1 and 3 was calculated for each sequence (Supplementary Scripts 1 and 2). Strong consensus sequences were identified for each selection, with the most abundant peptide sequences P28S1 and P29S1 at 47% and 62% occurrence for the P28 and P29 selections, respectively (Figure 3B and Figures S4-S5). The peptides also showed strong enrichment between the first and second round of selection, with P28S1 and P29S1 showing 586- and 750-fold enrichment, respectively (Figures S4-S5). The three most abundant peptides from each selection were synthesized and characterized by LC and ESI-MS (Figures S10-S15, Table S1). Their binding to the Spike RBD was then measured using biolayer interferometry (BLI). All peptides showed mid to low micromolar affinity for a commercially purchased Spike-RBD-Fc fusion protein (Figure 3B, Figure S6).

**Figure 3:**
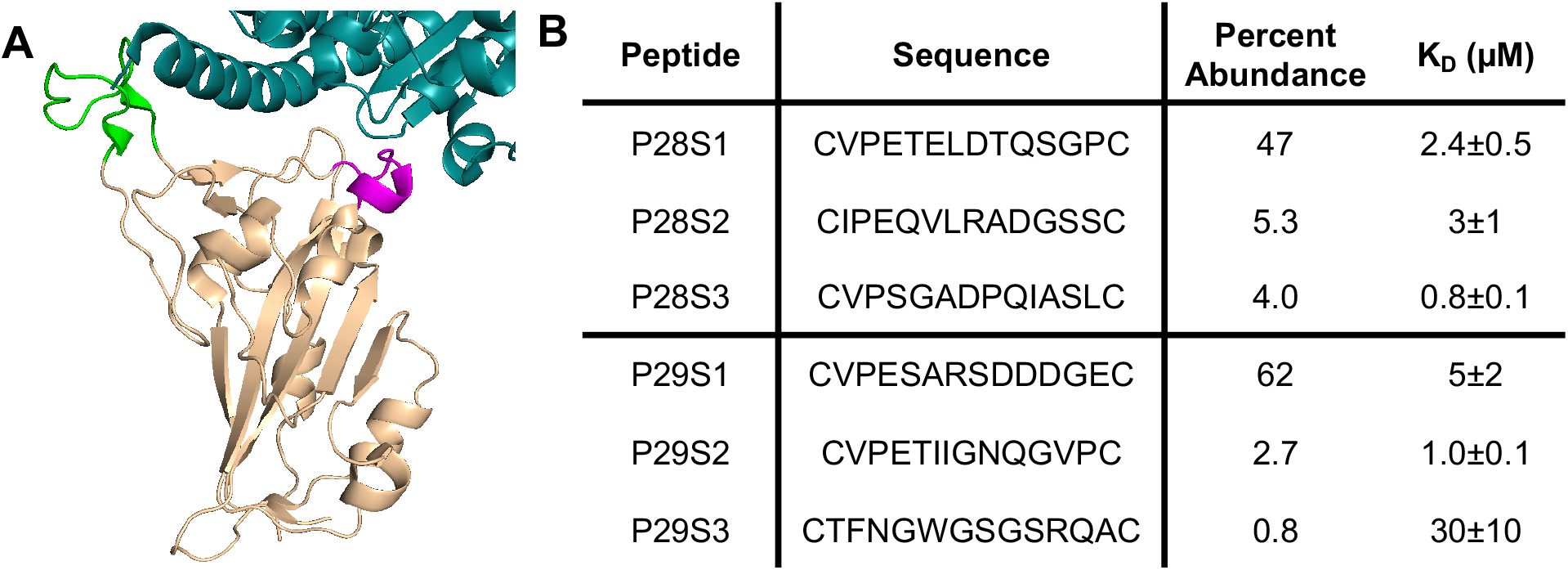
Selection of macrocyclic peptide ligands for the SARS-CoV-2 Spike RBD. **A)** Loops of the Spike RBD (beige) involved in recognition of ACE2 (Cyan) were identified for screening. P28 corresponds to the green residues and P29 corresponds to the magenta residues. **B)** The most abundant peptides from the selections against P28 and P29 are shown. Percent abundance corresponds to the percent of each sequence after the third round of selection against the corresponding peptides. Each peptide was analyzed for binding to the Spike RBD using BLI and the reported K_D_ values are given as mean ± SD of three independent experiments.

Encouraged by the BLI results, we proceeded to test inhibition of the Spike:ACE2 interactions by these identified macrocyclic peptides using a previously developed AlphaLISA assay.^35^ Excitingly, the top peptides displayed inhibition of the Spike:ACE2 interactions, with IC_50_ values of 1.5 ± 0.6 μM and 0.3 ± 0.1 μM for P28S1 and P29S1, respectively (Figures 4A, 4B and 4D; Figure S7). In spite of showing binding to the Spike RBD, less-enriched peptides showed low inhibition or no inhibition of the Spike:ACE2 interactions (Figure S7). This may be due to interactions with conformers of the Spike RBD that do not bind ACE2, as the peptides that we selected for were more than likely composed of binders for a variety of conformations of P28 and P29. Following the promising biochemical results of the highly enriched peptides, the peptides were then tested for viral inhibition of SARS-CoV-2 replication using a cytopathogenic effect (CPE) assay in Vero E6 cells. While P28S1 gave no detectable inhibition, P29S1 inhibited viral replication with an EC_50_ value of 3.1 ± 0.4 μM (Figures 4A and 4C). To our knowledge, this is the first *de novo* peptide that has exhibited complete viral inhibition by disrupting the interactions between the RBD and ACE2. We hypothesize that the discrepancy in the IC_50_ results and the CPE assay may be due to the orientation of the selected regions in the trimer of the Spike protein.^33^ With P28, the loop is oriented away from the center of the protein, so to prevent interactions with ACE2 at this region it may be necessary to block all three monomers at once. P29 loops, however, are closely packed in the center of the trimer, so binding to this region may be more efficient at blocking interactions between Spike and ACE2. Nevertheless, future structural experiments would give more insight into the interactions between P29S1 and the trimeric Spike protein. Also, the structures studies could be used to improve the binding affinity of P29S1, and it may be interesting to take advantage of a possible chelation effect by combining P29S1 and P28S1.

**Figure 4:**
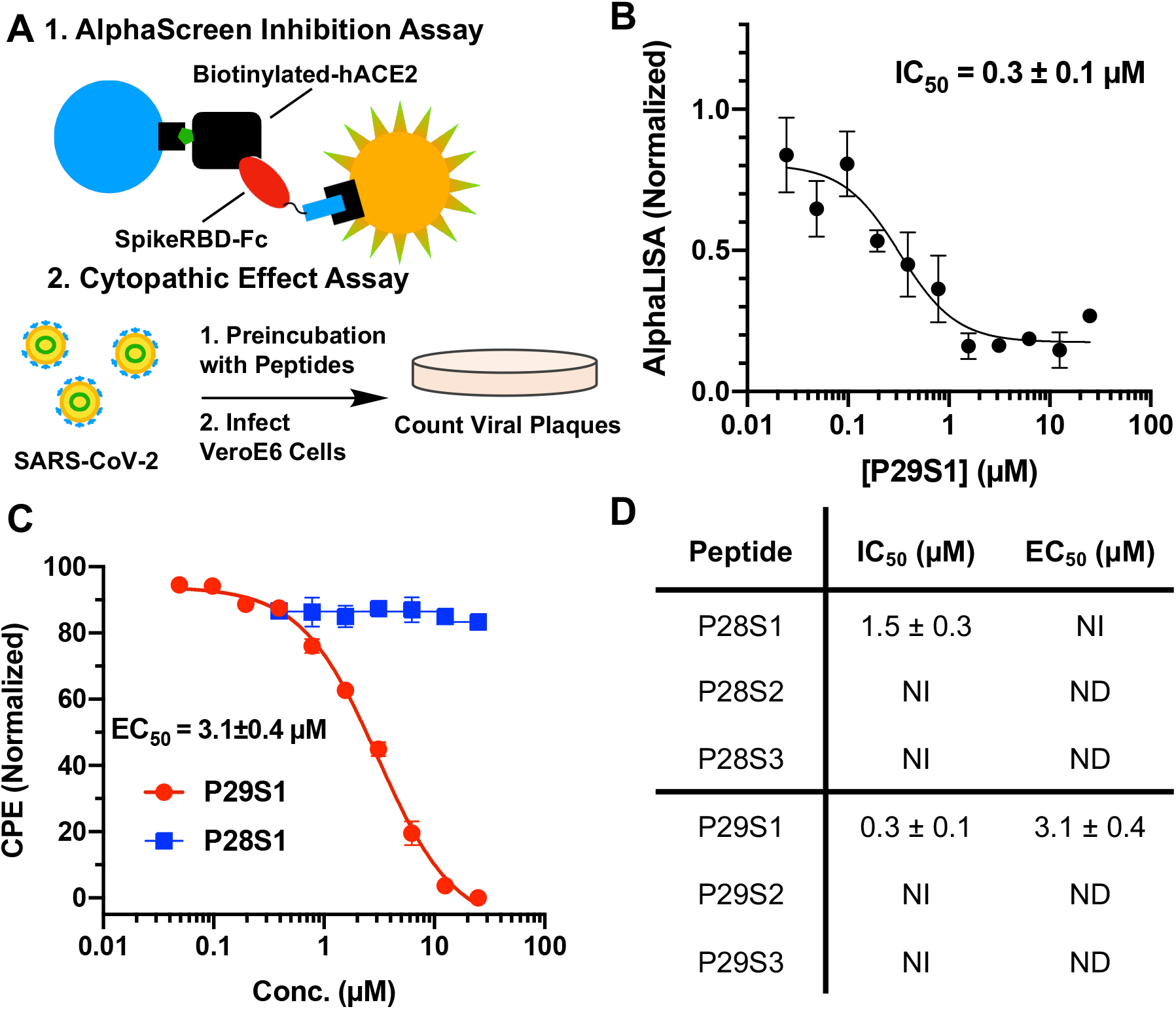
Characterization of macrocyclic peptides for inhibition of interactions between the Spike RBD and ACE2. **A)** Assays used to characterize inhibition consisted of an AlphaScreen assay to measure protein-protein interactions along with a cytopathogenic effect (CPE) assay that quantified infectivity of SARS-CoV-2 in Vero E6 cells. **B)** Inhibition of Spike RBD:ACE2 interactions by P29S1 using the AlphaScreen assay. Data points are presented as mean ± SEM of four biologically independent replicates (N=3). **C)** Live virus inhibition using P29S1 (red) and P28S1 (blue). Data points are presented as mean ± SD of three biologically independent experiments (N = 3). **D)** Summary of inhibition data for each of the tested peptides. NI = No inhibition. ND = Not determined. All data is reported as mean ± SD for three independent experiments (N = 3).

## CONCLUSION

In summary, we report the development of CAmCBT as a novel asymmetric macrocyclic peptide linker for phage display and its use in the construction of a phage-displayed macrocyclic peptide library for the identification of a potent neutralizing peptide P29S1 for SARS-CoV-2. CAmCBT shows quick condensation selectively with the *N*-terminal cysteine in a peptide/protein that completes within 3 hours at room temperature. Because of the asymmetric nature of CAmCBT, it can be further modified to include additional chemical functionalities for the introduction of other chemical fragments for expanding the chemical diversity and organic reactivities of phage-displayed peptides. We are currently exploring this potential. It is important to note that while the CAmCBT linker was in development, *Zheng et al*. reported an alternative strategy to cyclize a peptide with a *N*-terminal cysteine and an internal cysteine.^36^ In comparison to the linker reported in Zheng *et al*., CAmCBT reacts faster and is more synthetically accessible.

While we have observed promising antiviral activity of P29S1 through the inhibition of the interactions between the SARS-CoV-2 Spike protein and its human host receptor ACE2, this could be further optimized through several different strategies. As the protein exists as a trimer in the virion, it may be beneficial to develop a homotrimer to further improve the binding of P29S1 to the Spike protein. Additional structure-activity relationship studies may lead to more potent inhibitors with the guide of the structural analysis of P29S1 binding to the Spike RBD. A number of current SARS-CoV-2 variants have mutations in P28 and P29 epitopes. The reported method might also be quickly diverted to select macrocyclic peptides that neutralize these new viral variants. This possibility will be explored. Another potential benefit of using epitopes instead of full-length proteins for selection is the likely use of D-peptide epitopes for the selection of L-macrocyclic peptides that can be further converted to D-macrocyclic peptides for targeting the native L-form proteins. Due to their high metabolic stability and low immunogenicity, D-macrocyclic peptides are comparably more likely turned to be therapeutics than L-macrocyclic peptides. This potential will be explored as well to identify additional SARS-CoV-2 neutralizing macrocyclic peptides.

## METHODS

### 1. Primer Information and Plasmid Construction

#### Primer List

1. TEV-CAC_5_-sfGFP-F: 5’CGGCCGCGGCCTGCGTTAGCAAAGGTGAAGAACTG3’
2. TEV-CAC_5_-sfGFP-R: 5’CGGCGCACTGAAAATACAGGTTTTCCATGGTTAATTCCTCCTG3’
3. 12mer-pIII-NcoI-F: 5’CGGCCATGGCCTGC(NNK)_12_TGCGCGGCGAAAGCGGCCGGCCC3’
4. pIII-NcoI-R: 5’CATGCCATGGCCGGCTGGGCCGC3’
5. 5mer-pIII-NcoI-F: 5’CGGCCATGGCCTGC(NNN)_5_TGCGCGGCGAAAGCGGCCGGCCC3’
6. NGS-F1: 5’TCGTCGGCAGCGTCAGATGTGTATAAGAGACAGGCCCAGCCGGCCATG3’
7. NGS-R1: 5’GTCTCGTGGGCTCGGAGATGTGTATAAGAGACAGCGGCCGCTTTCGCCGC3’
8. NGS-i7: 5’CAAGCAGAAGACGGCATACGAGAT[i7]GTCTCGTGGGCTCGG3’
9. NGS-i5: 5’AATGATACGGCGACCACCGAGATCTACAC[i5]TCGTCGGCAGCGTC3’

#### Plasmid Construction

##### pADL-(NNK)_12_-gIII

The plasmid pADL-10b (Antibody Design Labs, San Diego, CA) was amplified with Phusion high-fidelity DNA polymerase using the primers 12mer-pIII-NcoI-F and pIII-NcoI-R. The product was digested with NcoI, ligated overnight with T4 DNA ligase, and used to transform electrocompetent *E. coli* Top10 (1 ng DNA per μL of cells). Transformed cells were diluted 1:9 in 2xYT and incubated at 37°C. After 1 hour, 100 μL was removed for to quantify transformation efficiency and the remaining culture was diluted to OD_600_ < 0.2 in fresh 2xYT media, supplemented with 100 μg·μL-1 ampicillin, and grown at 37°C overnight. In total 1.1 × 10^9^ transformants were obtained. The following day, the plasmid was extracted using a commercial extraction kit (Qiagen) and used to transform electrocompetent E. coli ER2738 exactly as described above yielding 1.8 × 10^9^ transformants.

##### pADL-(NNN)_5_-gIII

The plasmid pADL-10b (Antibody Design Labs, San Diego, CA) was amplified with Phusion high-fidelity DNA polymerase using the primers 5mer-pIII-NcoI-F and pIII-NcoI-R. The plasmid product was digested, ligated, and used to transform *E. coli* as described above to yield 2.0 × 10^9^ total transformants.

##### pBAD-TEV-CA_5_C-sfGFP-His_6_

The plasmid pBAD-sfGFP has been described previously.^37^ To introduce the polyalanine model peptide and TEV protease recognition sequence to the N-terminus of sfGFP, the plasmid pBAD-sfGFP was PCR amplified with primers TEV-CA_5_C-sfGFP-F and TEV-CA_5_C-sfGFP-R using Phusion high-fidelity DNA polymerase. The PCR product was phosphorylated with T4 polynucleotide kinase, followed by ligation with T4 DNA Ligase, and used to transform *E. coli* Top10 cells.

### 2. Biological Methods

#### Expression of CA_5_C-sfGFP

*E. coli* (Top10) containing the plasmid pBAD-TEV-CA_5_C-sfGFP-His_6_ were grown at 37 °C in 1 L of 2xYT containing 100 μg/mL ampicillin to OD_600_ = 0.6. Protein expression was induced by addition of 0.2% arabinose at 37 °C for 18 h. Cell pellets were then harvested by centrifugation (3750 xg, 20 min) and cells were resuspended in 40 mL of resuspension buffer (50 mM NaH_2_PO_4_, 300 mM NaCl, 10 mM imidazole, pH 8.0) and lysed by sonication. The lysate was clarified (10,000 xg, 30 min), and the supernatant incubated with 2 mL of Ni-NTA resin for 1 hour at 4 °C. The resin was then washed with 20 mL of resuspension buffer before eluting the protein with elution buffer (resuspension buffer + 250 mM imidazole). The eluant was concentrated and buffer exchanged to TEV protease buffer (50 mM Tris, 150 mM NaCl, 0.5 mM EDTA, pH 8.0) by centrifugation (MWCO 10 kDa). The protein was then incubated with TEV protease (1:50 enzyme:substrate ratio) for 48 hours at 4 °C. Complete digestion was confirmed through mass spectrometry. Following digestion with protease, the protein was concentrated to 1.4 mg/mL in storage buffer (TEV protease buffer + 20% glycerol) at -80 °C until further use. Protein concentration was determined by measuring the absorbance of the GFP chromophore at 485 nm using the reported extinction coefficient of 83,300 M^-1^·cm^-1^.

#### Cyclization of CA_5_C-sfGFP

Purified CA_5_C-sfGFP was diluted to 500 nM in PBS (pH 7.4) containing 1 mM TCEP and left to incubate for 30 min at 37 °C. Linker (100 μM final concentration) or equivalent amounts of DMSO were then added to the solution and it reacted for 3 hours at room temperature. The reaction mixtures were then concentrated and exchanged to ammonium biocarbonate (50 mM, pH 8.0) using an Amicon centrifugal filter (MWCO: 10 kDa). Samples were then analyzed using a QExactive Orbitrap LC-MS System (Thermo Fisher).

#### Phage Expression and Purification

*E. coli* ER2738 containing the pADL-NNK_12_-gIII or pADL-NNN_5_-gIII phagemid library were grown at 37°C in 250 mL of 2xYT media supplemented with 100 μg/mL ampicillin, 10 μg/mL tetracycline, and 1% glycerol. Upon reaching OD_600_ = 0.5-0.6, 20 mL of the culture of was transferred to a small flask and infected with 20μL of CM13d3 helper phage (Antibody Design Labs, San Diego, CA) at 37°C with shaking. After 45 min the cells were pelleted (3750 x *g*, 15 min) and resuspended in 200 mL of 2xYT media containing 10 μg/mL tetracycline, 100 μg/mL ampicillin, 25 μg/mL kanamycin, and 1 mM IPTG and incubated at 30 °C. 18 h post induction, the culture was transferred to 50 mL tubes, cells were pelleted (3750 x *g*, 20 min), and the supernatant was decanted into new tubes. Phages were precipitated by addition of appropriate amounts of 5x Precipitation Buffer (20% polyethylene glycol 8000, 2.5 M NaCl) to afford a 1x solution, then incubated at 4 °C for 1.5 h. The solution was centrifuged at 10,000 x *g* for 30 minutes, then the supernatant was discarded and the pellet resuspended in Binding Buffer (10 mM HEPES, 150 mM NaCl, 10 mM MgCl_2_, 1 mM KCl, pH 7.4; 2 mL per 50 mL tube). The resuspended phages were combined to one tube, the phage precipitation was repeated, and phages were ultimately resuspended in 2 mL of Binding Buffer. Any residual bacteria were then pelleted (13,500 x *g*, 20 min) and the supernatant was transferred to a fresh tube. The solutions incubated at 65 °C for 15 min to kill any remaining bacteria before being stored at 4 °C until further use.

#### Phage Quantification

For all experiments, phages were quantified via a colony forming unit assay. In this assay, serial dilutions of the phage solution were prepared in 2xYT media and 10 μL of each dilution was added to 90 μL of log-phase *E. coli* ER2738. Following addition of the phage dilutions, the culture was incubated at 37 °C for 45 min and then 10 μL was spotted in triplicate onto agar selection plates containing either 100 μg/mL ampicillin and 10 μg/mL tetracycline, which were incubated at 37 °C overnight. The following day, colonies in each spot were counted and this number was used to calculate the number of colony forming units in the solution.

#### Labelling Phages with FITC-CBT

The purified phage library (5mer) or M13KO7 (Antibody Design Labs) in PBS (pH 7.4) was incubated with 1 mM TCEP for 30 min at 37 °C followed by the addition of 50 μM FITC-CBT. The reaction was incubated at room temperature for 2.5 h followed by precipitation of the proteins with the addition of trichloroacetic acid. The precipitated protein pellet was washed twice with 500 μL of ice-cold acetone and the fluorescence was visualized under blue light.

#### Phage Infectivity Assay

For determining the effect of treatment with CAmCBT on phage infectivity, the purified phagemid library in PBS (pH 7.4) was incubated under various reaction conditions including: (1) PBS only, (2) 1 mM TCEP for 30 min at 37°C followed by 3 h at room temperature, (3) 30 min at 37°C followed by the addition of 50 μM CAmCBT and incubation for 3 h at room temperature, and (4) 1 mM TCEP for 30 min at 37°C followed by the addition of 100 μM CAmCBT (or equivolume amounts of DMSO) and incubation for 3 h at room temperature. Following incubation under the various conditions, the reaction was quenched by precipitation of phages (see Phage Purification section) and phages were resuspended in PBS. Phage solutions were then titered using the previously described colony forming unit assay.

#### Phage Library Cyclization with CAmCBT

The purified phage library (10^11^-10^12^ cfu) in Binding Buffer was incubated with 1 mM TCEP at 37°C. After 30 min, CAmCBT (10 mM in DMSO, 1% DMSO final) was added to a final concentration of 100 μM. The solution was incubated for 3 h at room temperature with slow rotation. Following the reaction, any remaining TCEP was quenched through addition of 10 mM oxidized glutathione followed by incubation at room temperature for 30 min. The library was then directly used for affinity selection without further purification.

#### Affinity Selection Against Peptides P28 and P29

##### Phage Selection

Streptavidin coated magnetic beads (100 μL, 50% slurry, Cytiva Sera-Mag) were transferred to a 1.5 mL tube, washed three times with 1 mL of Binding Buffer (10 mM HEPES, 150 mM NaCl, 10 mM MgCl_2_, 1 mM KCl, pH 7.4), resuspended in 100 uL of Binding Buffer, and split into two tubes. 500 pmol of biotinylated peptide was added to 1 mL of Binding Buffer in one of the tubes and an equal volume of Binding Buffer was added to the other tube (hereafter referred to as + tube and – tube, respectively). The beads/peptide mixture was incubated with rocking for 15 min at room temperature. The supernatant was then removed and the beads were washed with Binding Buffer (3 × 1 mL). 0.25 mL of 5x blocking buffer (Binding Buffer + 5% BSA + 0.5% Tween 20) was added to each tube and phage solution, and they incubated at room temperature with end-over-end rotation for 30 min. The blocking buffer was removed from the – tube, and the purified phage library was incubated with the beads for 30 min at room temperature as a negative selection. The supernatant was then transferred to the resin of the + tube and left to incubate at room temperature for 30 min. After 30 min the supernatant was removed and the resin was washed with Wash Buffer (Binding Buffer + 0.1% Tween 20) to remove nonspecifically bound phages. For the first round of selection, 8 × 1 mL washes were performed, then 10 × 1 mL washes were done to increase stringency in the second and third rounds. During each washing step the resin was completely resuspended by pipetting up and down. To remove phages binding to the polypropylene tube, the resin was transferred to fresh tubes after every other wash. After the last wash, phages were eluted by incubating for 15 min with 100 μL of Elution Buffer (50 mM glycine, pH 2.2). The supernatant was then removed and immediately added to 50 μL of Neutralization Buffer (1 M Tris, pH 8.0). The neutralized elution was immediately used for amplification.

##### Phage Amplification

A small aliquot (10 μL) of the phage elution was removed for quantification of phages. The remaining solution was added to an actively growing culture of *E. coli* ER2738 (OD_600_ = 0.5-0.6) in 20 mL 2xYT containing 10 μg/mL tetracycline for 45 min at 37 °C with rotation. After 45 min the cells were pelleted (3750 x *g*, 15 min), resuspended in 200 mL of 2xYT containing 100 μg/mL ampicillin and 10 μg/mL tetracycline, and amplified overnight at 37 °C. The following day, phagemids were extracted from the amplified culture using a commercial plasmid purification kit. The remaining culture was then inoculated into 100 mL of fresh 2xYT containing 100 μg/mL ampicillin and 10 μg/mL tetracycline for subsequent phage expression.

#### Illumina Sequencing of Selected Phage Libraries

A four step PCR cycle was used to amplify the library region out of the original phagemid library using primers NGS-F1 and NGS-R1, as has been previously described.^34^zacx The amplicons were purified and extracted from a 3% agarose gel according to a GenCatch gel extraction kit, then indices were attached using a subsequent PCR with NGS-i7 and NGS-i5 primers. To identify the rounds of selection, each round contained a unique combination of i7 or i5 indices. The PCR products were purified using a GenCatch gel extraction kit and submitted to the Genomics and Bioinformatics center at Texas A&M University for sequencing on an Illumina iSeq (4M Reads, 2×150bp). Sequences were analyzed for enrichment in R, and all scripts are available in the Supplementary Information.

#### Biolayer Interferometry Assays

Biolayer interferometery (BLI) experiments were performed using AHC biosensors (Forte Bio) on an Octet Red 96 biolayer interferometer (Forte Bio). All experiments were performed in assay buffer (PBS + 0.05 mg/mL BSA). Fc-Spike-RBD (100 nM in Assay Buffer) was loaded onto the sensor to 1-2 nm of loading over 5 minutes, then the sensors were quenched for one minute with a 1% BSA solution in assay buffer. Peptides were serially diluted in assay buffer (75 - 9.4 μM) and the association and dissociation to Fc-Spike-RBD were measured over 2 and 5 minutes, respectively. Between each experiment, sensors were regenerated using 10 mM glycine, pH 1.5. Binding curves were referenced against a protein-only sensor and screened for nonspecific binding prior to analysis. All curves were fit in the Octet Red Kinetic Analysis program using a 1:1 protein:ligand binding model.

#### AlphaScreen Assays

AlphaScreen assays were performed similar to a previously reported protocol for Spike:ACE2 interactions (CITE). Final concentrations in the assays are as follows: 4 nM Spike-RBD, 4 nM ACE2, 10 μg/mL protein A acceptor beads, 10 μg/mL streptavidin donor beads, and varying concentrations of inhibitor. All proteins were prepared in assay buffer (PBS + 0.05 mg/mL BSA). The peptides were serially diluted in assay buffer (PBS + 0.05 mg/mL BSA) that contained 0.16% TEA to appropriate concentrations and were preincubated with Fc-Spike-RBD (R&D Systems, Cat. No. 10542-CV-100) at 37 °C for 1 hour (12.5 μL total volume). These were then added to AlphaScreen plates (Perkin Elmer, 384-well) that had been preincubated for 30 minutes with 12.5 μL of Biotinylated-ACE2 (Sino Biological, Cat. No. 10108-H27B-B). After a 30 minute incubation at 37 °C, 25 μL of premixed acceptor/donor beads were then added to the plates and incubated for 30 minutes at 37 °C. After incubation, the Alpha signal was read using a Neo2 plate reader (Biotek). Data was then normalized, with 100% being the positive control (no peptides added) and 0% being just ACE2 added. The normalized data was plotted and fitted using a nonlinear regression model (4 parameters) on GraphPad Prism.

#### SARS-CoV-2 Cytopathic Effect Assay

Vero cells (ATCC-CCL81) were maintained in M199 media supplemented with 5% fetal bovine serum and 1% penicillin-streptomycin-antimycotic. All cells were maintained at 37 °C and 5% CO_2_ in a humidified incubator. P29S1 and P28S1 dilutions were prepared in MilliQ water containing 2% triethylamine, along with a blank series containing 2% triethylamine in water. The solutions were mixed 1:1 with SARS-CoV-2 (SARS-CoV-2, 2019-nCoV/USA-IL1/2020) in M199 cell culture media and incubated at 37 °C for 1 hour. After incubating on cells, a 1% agarose overlay was added to each well. After 24 hours, a secondary 1% agarose overlay containing neutral red was added to each well. After another 24 hours, viral plaques were counted. Inhibition for each peptide was then calculated by normalizing the plaques with the total plaques formed in the 2% triethylamine blank series (blank = 100, no plaques = 0).

### 3. Synthetic Methods

#### Synthesis of CAmCBT

The synthesis of 6-amino-1,3-benzothiazole-2-carbonitrile (**4**) was performed and compounds were characterized using NMR as previously described.^38^ Detailed procedures are given below.

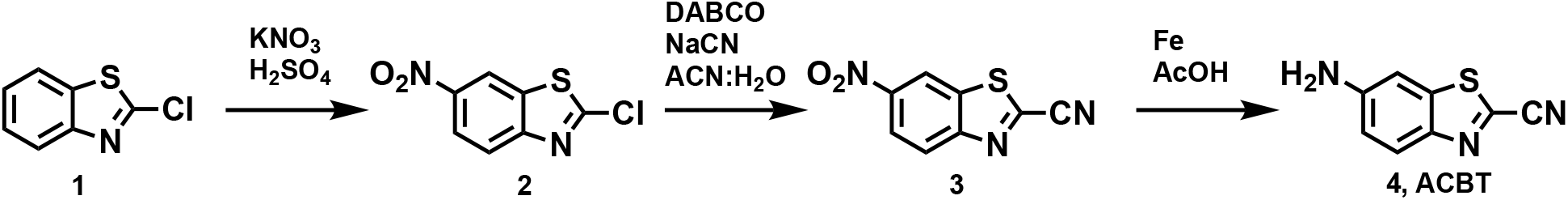

2-Chloro-6-nitro-1,3-benzothiazole (**2**)

2-Chloro-1,3-benzothiazole (10.0 g, 58.95 mmol) was added drop-wise to 60 mL of H_2_SO_4_ that had been chilled to 4 °C. Potassium nitrate (6.56 g, 64.85 mmol) was then added and the solution was left to stir at 4 °C for 30 minutes, then reacted at room temperature for 1 hour. The reaction mixture was then poured onto ice and the precipitate was collected and rinsed with cold water. The crude product was recrystallized using 600 mL of ethanol, giving **2** as yellow needles. The product was used for the next step without further purification.

6-Nitro-benzo[*d*]thiazole-2-carbonitrile (**3**)

Sodium cyanide (1.470 g, 29.99 mmol) was dissolved in 60 mL of H_2_O and slowly added to a stirring solution of **2** (6.14 g, 28.61 mmol) and DABCO (450 mg, 4.21 mmol) in 600 mL of MeCN. The reaction mixture was then stirred for 24 hours at room temperature. Remaining cyanide was quenched through the addition of 20 mL of 0.3 M FeCl_3_ and 300 mL of H_2_O were added to the flask. The organic layer was extracted using ethyl acetate (3 × 250 mL) and then washed with 80 mL of brine. The product was dried with Na_2_SO_4_, filtered, and concentrated in vacuo to give a brown solid. The crude product was then loaded onto a silica plug, eluted with 500 mL of chloroform, and concentrated in vacuo. Product **3** was then used without further purification (4.37 g, 74%).

6-Amino-benzo[*d*]thiazole-2-carbonitrile (**4**, ACBT)

Compound **3** (4.37 g, 21.30 mmol) was dissolved in 500 mL of acetic acid. Iron powder (61.43 g, 1099.51 mmol) was added to the flask and the reaction mixture stirred at room temperature for 24 hours. The product was diluted with water (900 mL) and filtered through celite. The solution was then extracted using ethyl acetate (4 × 400 mL) and the organic layers were combined, washed with brine (2 × 250 mL), then concentrated in vacuo. The crude product was loaded onto a silica plug, flushed with 1 L of chloroform, and concentrated in vacuo to afford **4** as an orange solid.

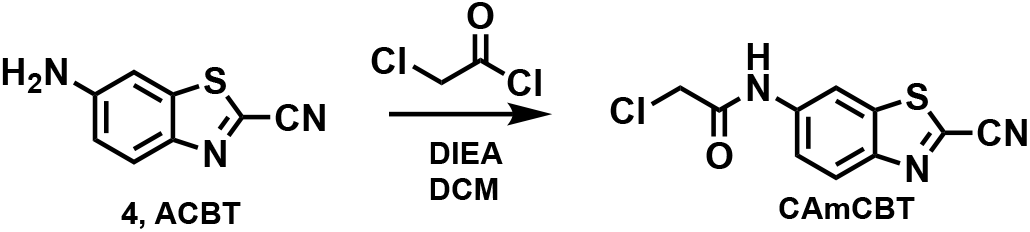

2-Chloro-*N*-(2-cyanobenzo[*d*]thiazol-6-yl)acetamide (**CAmCBT)**

Disopropylethylamine (0.252 mL, 1.37 mmol) and chloroacetyl chloride (0.06 mL, 0.685 mmol) was added to a solution of **4** (0.120 g, 0.685 mmol) in dry DCM (5 mL) at 4 °C. The reaction was then left to stir at room temperature for 16 hours and the solvent removed in vacuo. The residue was then purified by silica gel column chromatography (10-70% ethyl acetate in hexane, 20 min) to afford **CAmCBT** (0.115 g, 59%) as a yellow solid.

#### Synthesis of FITC-CBT

Synthesis of FITC-CBT was done and the compound was characterized as previously reported.^39^ Detailed procedures are given below.

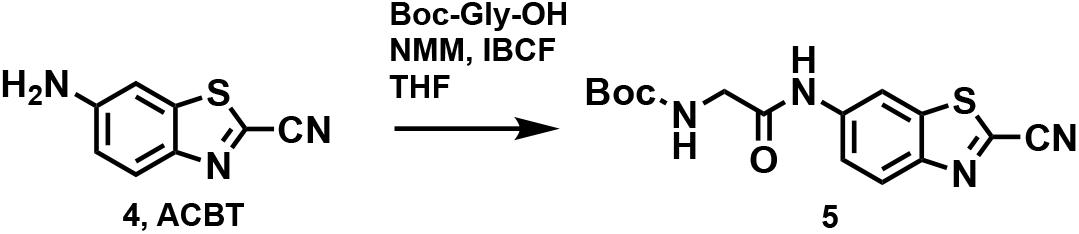

2-Amino-*N*-(2-cyanobenzo[*d*]thiazol-6-yl)acetamide (**5**)

Isobutylchloroformate (0.046 mL, 0.355 mmol), Boc-glycine (60 mg, 0.342 mmol), and *N*-methylmorpholine (0.038 mL, 0.305 mmol) were dissolved in THF at 4 °C and mixed for 20 minutes. Compound **4** (0.05 g, 0.286 mmol) was added to the reaction mixture and stirred overnight at room temperature. The reaction was quenched with sodium biocarbonate (saturated, 40 mL). Then the organic layer was extracted using ethyl acetate (2 × 50 mL) and the combined extracts were dried with magnesium sulfate. The crude product was purified using silica gel column chromatography, then the boc was deprotected by incubation with 20% TFA in DCM for 1 hour. The crude product was precipitated in ether to isolate compound **5** and directly used for the next step in synthesis.

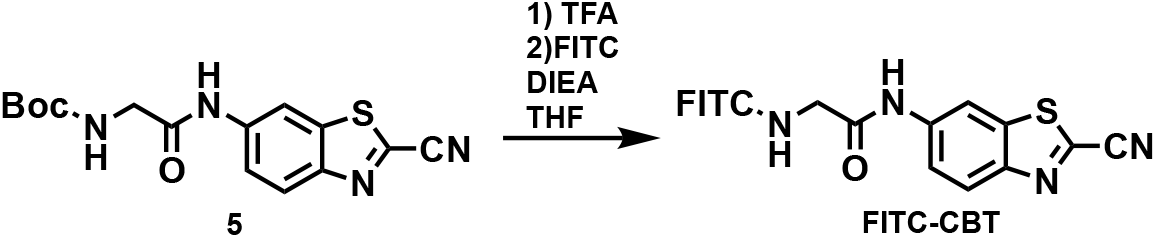

*N*-(2-cyanobenzo[*d*]thiazol-6-yl)-2-(2-(3’,6’-dihydroxy-3-oxo-3*H*-spiro[isobenzofuran-1,9’-xanthen]-5-yl)-2-thioxomethylidene-21^4^-diazaneyl)acetamide (**FITC-CBT**)

The amine **5** (50 mg, 0.18 mmol) was dissolved in a solution of diisopropylethyl amine (41 μL, 0.22 mmol) in dry methanol and THF (2:1, 1.2 mL). The solution was cooled to 0°C before adding fluorescein-5-isothiocyanate (FITC, 72 mg, 0.18 mmol). The reaction mixture was warmed to room temperature and stirred overnight before concentrating and purifying by column chromatography on silica gel using a gradient of DCM:acetone:MeOH (80:10:10 to 0:90:10) to afford **FITC-CBT** (71 mg, 62%) as an orange solid.

#### Synthesis of Fmoc-Aluc

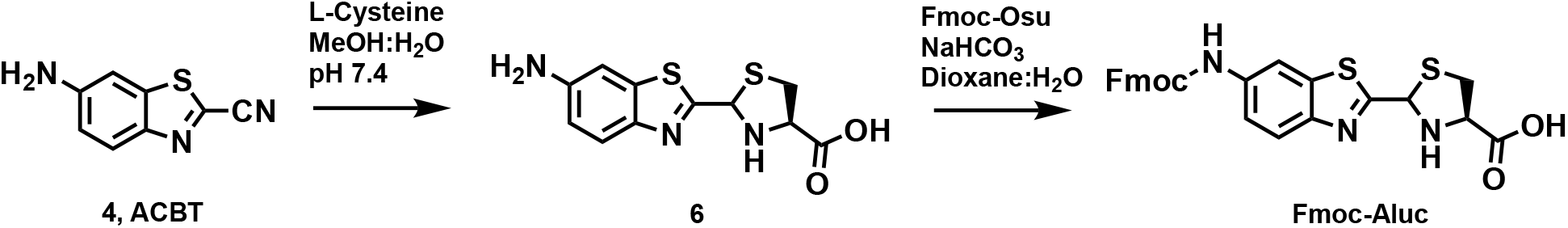

2-(6-aminobenzo[*d*]thiazol-2-yl)-4,5-dihydrothiazole-4-carboxylic acid (**6**, L-aminoluciferin) L-cysteine (432 mg, 3.56 mmol) was dissolved in 50 mL of water, and the pH was adjusted to 7.4 by addition of 1 M Na_2_CO_3_. Compound **4** (505 mg, 2.86 mmol) was dissolved in 20 mL of methanol and added to the reaction mixture. After reacting for 2.5 hours at room temperature, the methanol was evaporated under vacuum, then the pH was adjusted to 3 using 10% HCl, resulting in an orange precipitate. The precipitate was then filtered, lyophilized, and the crude material (**6**, 450 mg, 56% yield) was directly used for the next step in the synthesis.

2-(6-((((9*H*-fluoren-9-yl)methoxy)carbonyl)amino)benzo[*d*]thiazol-2-yl)-4,5-dihydrothiazole-4-carboxylic acid (**Fmoc-Aluc**)

Compound **6** (300 mg, 1.03 mmol) was dissolved in 10 mL of 1:1 Dioxane:H_2_O and Fmoc-Osu (420 mg, 1.24 mmol) was added and stirred at room temperature for 6 hours. Aqueous HCl (1 M, 10 mL) was added to the mixture and the organic layer was extracted with EtoAc (3 × 5 mL). The combined organic layers were washed with 1M HCl, water, and brine. Then the combined aqueous layers were extracted with EtOAc, dried and evaporated under reduced pressure. Crude product was then purified using flash column chromatography affording **Fmoc-Aluc** as a white solide (250 mg, 46% yield. ^1^H NMR (400 MHz, CDCl_3_) δ 8.31 (s, 1H), 8.06 (d, *J* = 8.9 Hz, 1H), 7.87 – 7.79 (m, 2H), 7.65 (d, *J* = 7.5 Hz, 2H), 7.46 (t, *J* = 7.7 Hz, 2H), 7.37 (td, *J* = 7.4, 1.2 Hz, 2H), 6.88 (s, 1H), 5.47 (t, *J* = 9.8 Hz, 1H), 4.66 (d, *J* = 6.3 Hz, 2H), 4.32 (t, *J* = 6.3 Hz, 1H), 3.85 (s, 1H), 3.82 (s, 1H).

#### Peptide Synthesis

Initial sequences of peptides (until N-terminal cysteine) were synthesized on a low loading ProTide rink amide resin (CEM #R002) using an automated Liberty Blue Peptide Synthesizer with an HT12 attachment. To prevent aspartimide formation, Fmoc-Asp(OtBu)-(Dmb)Gly-OH was used for synthesis of peptides containing DG residues. All other derivatives were standard derivatives for Fmoc peptide synthesis. Fmoc-amino acids were deprotected using 20% piperidine in DMF with a two-step microwave cycle: 1) 75 °C, 15 s 2) 90 °C, 50 s. Residues were then coupled with 0.125 M DIC and 0.25 M Oxyma using CEM’s standard microwave coupling cycle: 1) 75 °C, 15 s 2) 90 °C, 110 s. Following synthesis of peptides on the Liberty Blue, the linker was coupled by hand in a two step process. First, Fmoc-Aluc (15 mg, 1.2 eq) was coupled to the N-terminus using HATU (152 mg, 4 eq) and DIPEA (72 μL, 4 eq) in DMF (1 mL) and reacted for 16 hours at room temperature. Following this, the chloroacetyl moiety was added. Chloroacetyl chloride (8 μL, 4 eq) was added to the resin at 4 °C containing DIPEA (72 μL, 4 eq) and incubated for 3 hours at room temperature. The chloroacetyl chloride coupling was repeated once before cleavage. The peptides were cleaved from the resin by agitating for 3 hours in 3 mL of 92.5:2.5:2.5:2.5 TFA:H2O:DODT:TIS. The products were then filtered and peptides were precipitated out of the filtrate using cold ether. The precipitates were collected by centrifugation and washed again with cold ether, then lyophilized. The crude products were then dissolved in 1 mL of DMF containing 4 equivalents of DIPEA and cyclized for two hours at 42 °C. The peptides were purified by reverse-phase HPLC (Acetonitrile in Water with 0.1% formic acid) using either a Discovery BIO wide pore C18-5 column (25 cm x 10 mm, Millipore-Sigma #568230-U) or a Shim-pack GIS C18 (20 × 250 mm, 10 μm, Shimadzu). Following purification, fractions were analyzed using high resolution mass spectrometry. Corresponding masses for synthesized peptides are listed in Supplementary Table 1.

## ABBREVIATION

S: Spike
RBD: receptor binding domain
ACE2: angiotensin converting enzyme 2
ORF: open reading frame
CBT: cyanobenzothiazole
CAmCBT: *N*-chloroacetyl 2-chloro-*N*-(2-cyanobenzo[*d*]thiazol-6-yl)acetamide
eGFP sfGFP: superfolder green fluorescent protein
TEV: tobacco etch virus
TCEP: triscarboxyethylphosphine

## ACKNOWLEDGMENTS

We thank Dr. Alfred Tuley for his valuable advice during the synthesis of CAmCBT. This work was supported by National Institutes of Health (Grant R35GM145351), National Science Foundation (Grant 2111297), Welch Foundation (Grant A-1715), and the Texas A&M University X-Grants mechanism. We kindly acknowledge supports provided by the Genomics and Bioinformatics Service from the Texas AgriLife Research.

