## Supporting Information for "A Novel Regioselective Approach to Cyclize Phage-Displayed Peptides in Combination with Epitope-Directed Selection to Identify a Potent Neutralizing Macrocyclic Peptide for SARS-CoV-2"

### SUPPLEMENTARY FIGURES

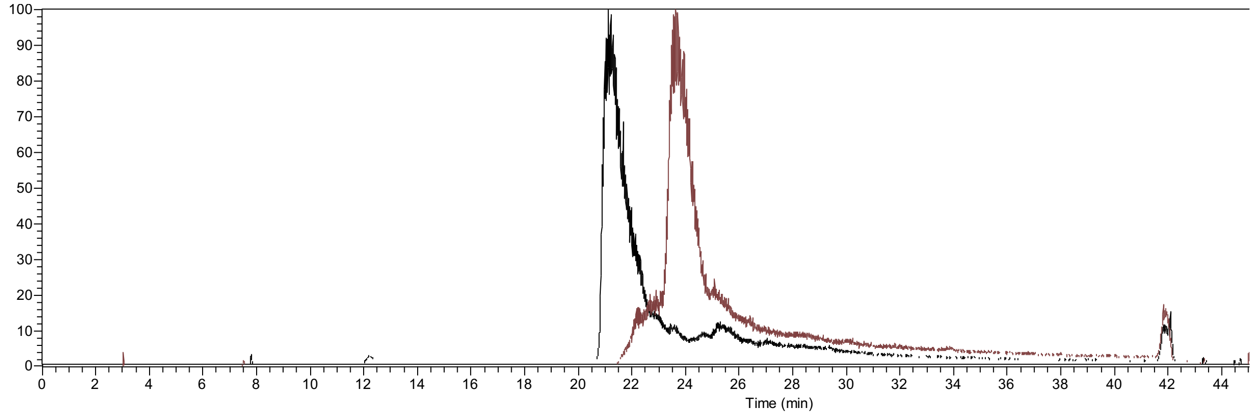


**Figure S1:** TIC chromatogram for ESI-MS of $CA_5C$-sfGFP before (black) and after (maroon) reaction with 100 μM of CAmCBT for 3 hours at room temperature.

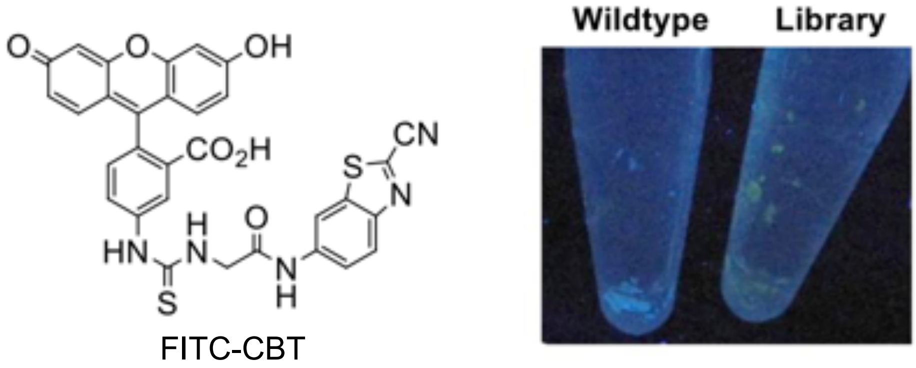


**Figure S2:** Labelling phages with FITC-CBT. Only the 5mer Library ($CX_5C$) containing an N-terminal cysteine showed fluorescence when reacted with FITC-CBT.

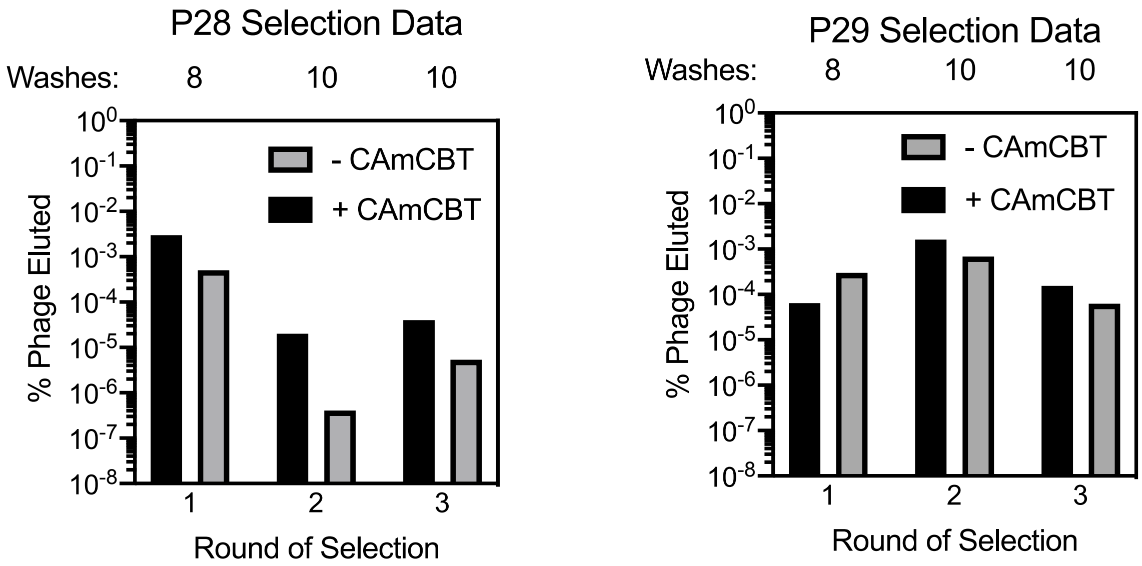


**Figure S3:** Phage titers during the selections against P28 and P29. Titers indicated enrichment of peptides binding when cyclized by CAmCBT.

**A**

p28_12mer_R3top100

| | V1 | V2 | V3 | V4 | V5 | V6 | V7 | V8 | V9 | V10 | V11 | V12 | V13 | V14 | n | percent |
|---|---|---|---|---|---|---|---|---|---|---|---|---|---|---|---|---|
| **P28S1** | C | V | P | E | T | E | L | D | T | TAG | S | G | P | C | 194735 | 47.4536331935404 |
| | C | V | P | E | S | A | R | S | D | D | D | G | E | C | 25557 | 6.22780960550139 |
| **P28S2** | C | I | P | E | Q | V | L | R | A | D | G | S | S | C | 22047 | 5.37248183951517 |
| **P28S3** | C | V | P | S | G | A | D | P | TAG | I | A | S | L | C | 16700 | 4.0695081743504 |
| | C | V | P | V | E | V | R | G | P | G | P | T | A | C | 14834 | 3.61479546456969 |
| | C | I | P | E | E | E | L | T | A | TAG | D | S | L | C | 9615 | 2.34301323930414 |
| | C | V | P | D | S | V | TAG | V | N | S | N | V | Q | C | 7676 | 1.87051166145591 |
| | C | V | D | P | F | V | Y | G | G | D | E | TAG | N | C | 2676 | 0.652096040392915 |
| | C | S | G | D | G | I | V | L | S | S | M | G | R | C | 2071 | 0.504667750244292 |
| | C | Q | I | V | W | D | H | P | TAG | N | S | T | L | C | 1884 | 0.459099006016536 |
| | C | I | P | P | E | A | N | G | L | TAG | G | W | R | C | 1636 | 0.398665591211812 |
| | C | I | P | TAG | A | A | D | P | N | S | W | D | L | C | 1503 | 0.366255735691536 |
| | C | V | P | E | A | V | R | G | D | V | G | S | S | C | 1331 | 0.324342238326969 |
| | C | V | P | E | S | E | L | N | S | L | P | R | P | C | 901 | 0.219558494915552 |

**B**

p28_12mer_R3enriched

| | V1 | V2 | V3 | V4 | V5 | V6 | V7 | V8 | V9 | V10 | V11 | V12 | V13 | V14 | n | enrichment |
|---|---|---|---|---|---|---|---|---|---|---|---|---|---|---|---|---|
| | C | S | G | D | G | I | V | L | S | S | M | G | R | C | 2071 | 1070.66953766001 |
| **P28S1** | C | V | P | E | T | E | L | D | T | TAG | S | G | P | C | 194735 | 586.571439597557 |
| | C | I | P | TAG | A | A | D | P | N | S | W | D | L | C | 1503 | 387.874774288506 |
| **P28S2** | C | I | P | E | Q | V | L | R | A | D | G | S | S | C | 22047 | 373.05068109633 |
| | C | V | P | V | E | V | R | G | P | G | P | T | A | C | 14834 | 273.145442533775 |
| | C | V | P | T | I | V | TAG | G | D | S | S | G | D | C | 252 | 259.802243834208 |
| **P28S3** | C | V | P | S | G | A | D | P | TAG | I | A | S | L | C | 16700 | 256.960049279274 |
| | C | V | P | E | A | V | R | G | D | V | G | S | S | C | 1331 | 228.581869406965 |
| | C | V | P | D | S | V | TAG | V | N | S | N | V | Q | C | 7676 | 219.669976154253 |
| | C | F | R | TAG | L | C | TAG | L | Y | M | A | L | S | C | 551 | 189.082058667524 |
| | C | V | P | E | T | E | L | D | T | L | S | G | P | C | 843 | 173.489120279553 |
| | C | V | P | L | S | V | TAG | A | Q | G | Y | T | A | C | 162 | 166.658585321991 |
| | C | V | P | E | TAG | V | I | S | E | D | T | T | S | C | 476 | 163.208820191909 |
| | C | V | P | E | S | I | Q | E | G | R | Q | TAG | M | C | 143 | 146.994924080523 |

**Figure S4:** Next-generation sequencing data for the selection against P28. Sequences were sorted by abundance (A) and enrichment between rounds 1 and 3 of selection (B). Enrichment was calculated by the (%R3-%R1)/%R1.

**A**

p29_12mer_R3top100

| | V1 | V2 | V3 | V4 | V5 | V6 | V7 | V8 | V9 | V10 | V11 | V12 | V13 | V14 | n | percent |
|---|---|---|---|---|---|---|---|---|---|---|---|---|---|---|---|---|
| **P29S1** | C | V | P | E | S | A | R | S | D | D | D | G | E | C | 179599 | 62.1916802581869 |
| **P29S2** | C | V | P | E | T | I | I | G | N | TAG | G | V | P | C | 8050 | 2.78756020956912 |
| | C | N | Y | S | A | TAG | P | I | P | C | V | L | M | C | 2722 | 0.942576259682876 |
| **P29S3** | C | T | F | N | G | W | G | S | G | S | R | TAG | A | C | 2311 | 0.800254862647732 |
| | C | S | G | I | S | D | C | L | F | V | T | A | R | C | 1683 | 0.582790538224203 |
| | C | L | F | L | S | I | A | I | V | S | T | P | S | C | 1314 | 0.455012933586811 |
| | C | Y | S | H | R | F | S | R | N | TAG | G | K | M | C | 689 | 0.238587451477407 |
| | C | F | R | L | F | R | F | Y | T | G | G | E | D | C | 525 | 0.181797404971899 |
| | C | V | P | E | S | V | R | S | D | D | D | G | E | C | 489 | 0.169331297202398 |
| | C | P | K | G | L | L | N | P | Y | F | H | N | Q | C | 425 | 0.147169327834395 |
| | C | F | K | S | R | TAG | G | V | G | R | R | G | V | C | 369 | 0.127777604637392 |
| | C | Y | I | TAG | V | G | D | N | G | H | TAG | F | N | C | 363 | 0.125699920009142 |
| | C | V | V | TAG | R | S | V | A | G | L | TAG | S | Y | C | 359 | 0.124314796923642 |
| | C | H | L | R | TAG | N | C | R | W | R | W | I | V | C | 316 | 0.109424723754515 |

**B**

p29_12mer_R3enriched

| | V1 | V2 | V3 | V4 | V5 | V6 | V7 | V8 | V9 | V10 | V11 | V12 | V13 | V14 | n | enrichment |
|---|---|---|---|---|---|---|---|---|---|---|---|---|---|---|---|---|
| **P29S1** | C | V | P | E | S | A | R | S | D | D | D | G | E | C | 179599 | 750.542033291433 |
| | C | V | P | E | S | V | R | S | D | D | D | G | E | C | 489 | 278.312821391841 |
| **P29S2** | C | V | P | E | T | I | I | G | N | TAG | G | V | P | C | 8050 | 247.545648837044 |
| | C | V | S | E | S | A | R | S | D | D | D | G | E | C | 215 | 244.612501428408 |
| | C | N | Y | S | A | TAG | P | I | P | C | V | L | M | C | 2722 | 171.753805914245 |
| | C | V | P | G | S | A | R | S | D | D | D | G | E | C | 106 | 120.092675122843 |
| **P29S3** | C | T | F | N | G | W | G | S | G | S | R | TAG | A | C | 2311 | 104.60195177694 |
| | C | V | P | E | S | A | R | S | D | D | D | G | G | C | 47 | 52.6920351959776 |
| | C | I | P | E | S | A | R | S | D | D | D | G | E | C | 35 | 38.9834304650897 |
| | Y | V | P | E | S | A | R | S | D | D | D | G | E | C | 32 | 35.5562792823677 |
| | C | F | P | E | S | A | R | S | D | D | D | G | E | C | 27 | 29.8443606444978 |
| | C | L | F | L | S | I | A | I | V | S | T | P | S | C | 1314 | 24.0182036338704 |
| | C | V | P | E | T | I | I | G | S | TAG | G | V | P | C | 17 | 18.4205233687579 |
| | C | G | S | TAG | F | G | V | V | V | V | Y | C | V | C | 291 | 17.4685369291129 |

**Figure S5:** Next-generation sequencing data for the selection against P29. Sequences were sorted by abundance (A) and enrichment between rounds 1 and 3 of selection (B). Enrichment was calculated by the (%R3-%R1)/%R1.

P28S1

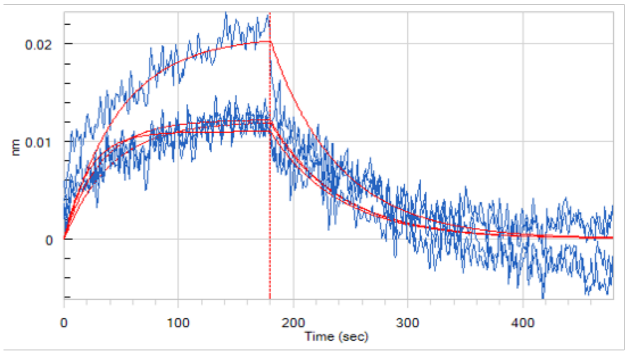


Conc. (µM): 6.25, 3.13, 1.56, 0.781
$K_D$: 2.4±0.5 µM

P29S1

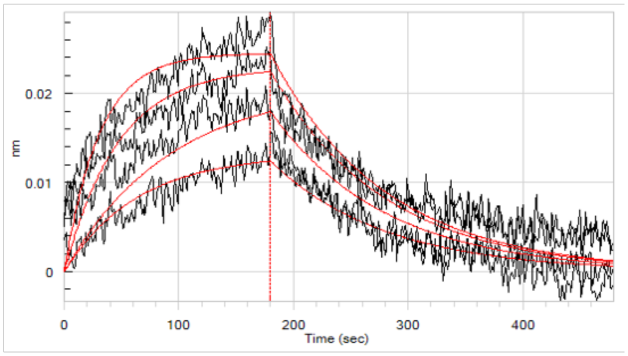


Conc. (µM): 25, 12.5, 6.25, 0.781
$K_D$: 5±2 µM

P28S2

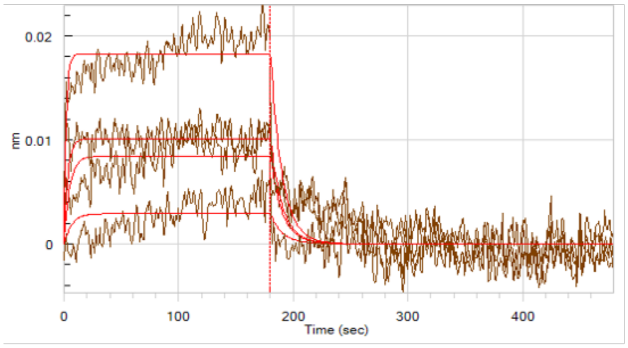


Conc. (µM): 12.5, 6.25, 3.13, 1.56
$K_D$: 3±1 µM

P29S2

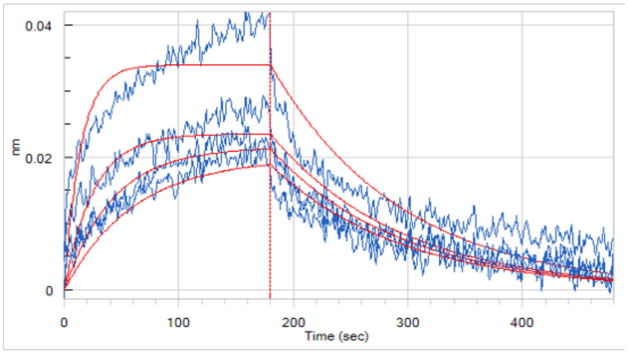


Conc. (µM): 6.25, 3.13, 1.56, 0.781
$K_D$: 1.0±0.1 µM

P28S3

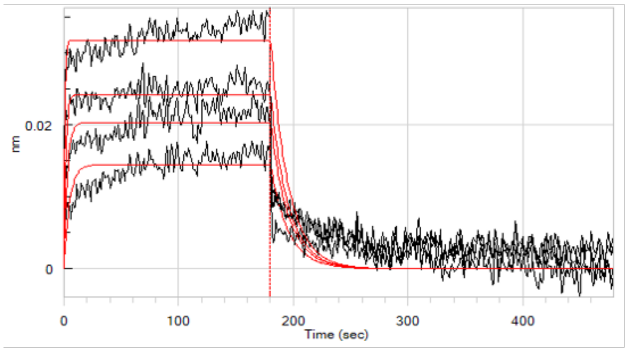


Conc. (µM): 12.5, 6.25, 3.13, 1.56
$K_D$: 0.8±0.1 µM

P29S3

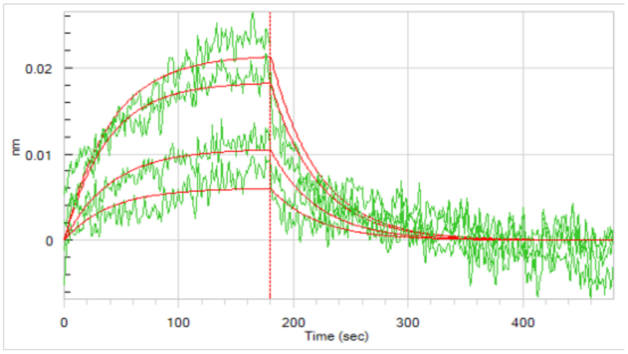


Conc. (µM): 6.25, 3.13, 1.56, 0.781
$K_D$: 30±10 µM

**Figure S6:** Biolayer interferometry (BLI) traces of the selected peptides against P28 and P29. Concentrations tested are listed below the traces for the corresponding peptides, along with $K_D$ values given as the mean ± s.d. of 3 independent experiments (n=3).

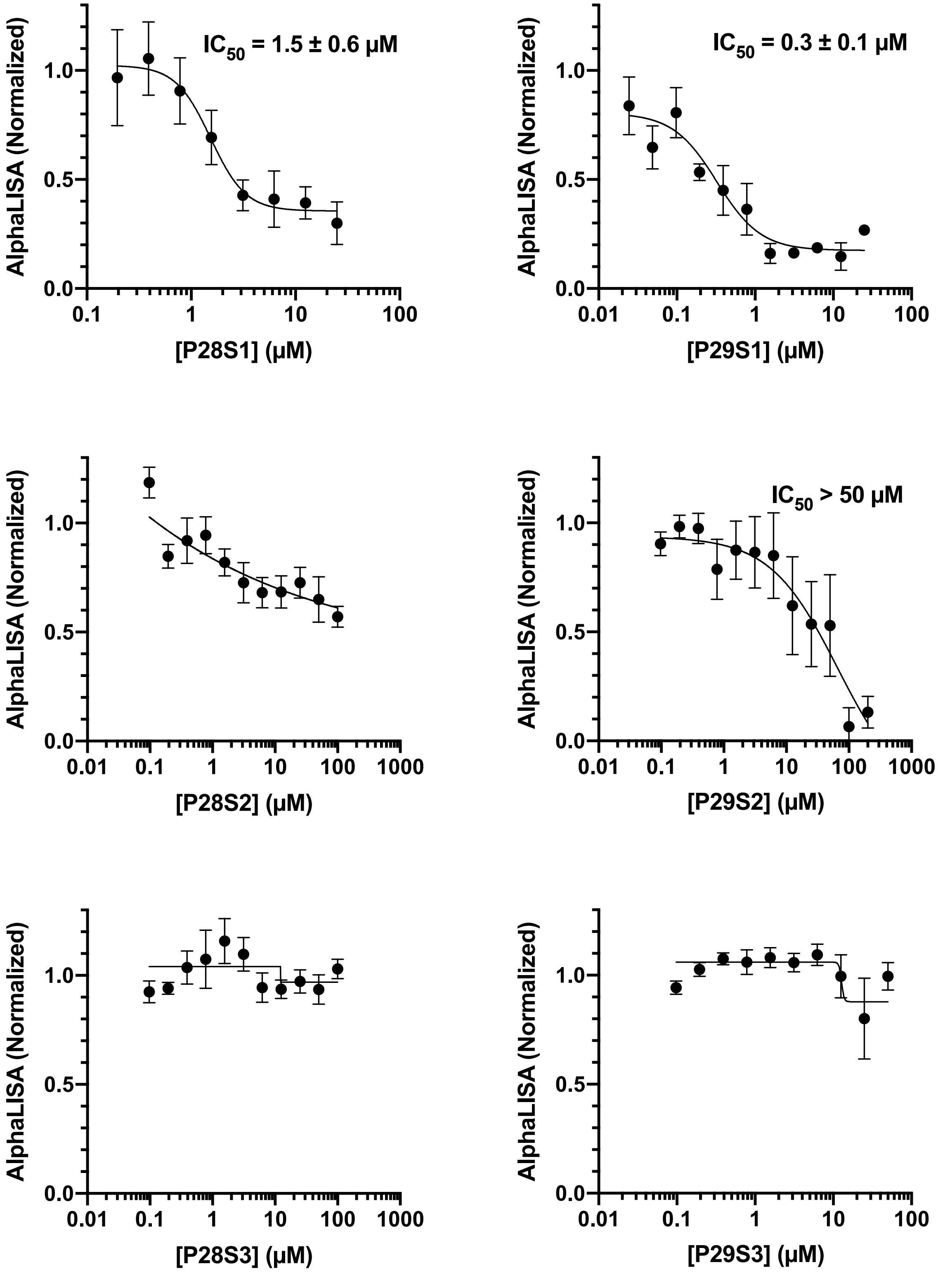


**Figure S7:** AlphaScreen data for the P28 and P29 selected peptides. Data points and $IC_{50}$ values are given as the mean ± s.d. of three biologically independent experiments (n = 3).

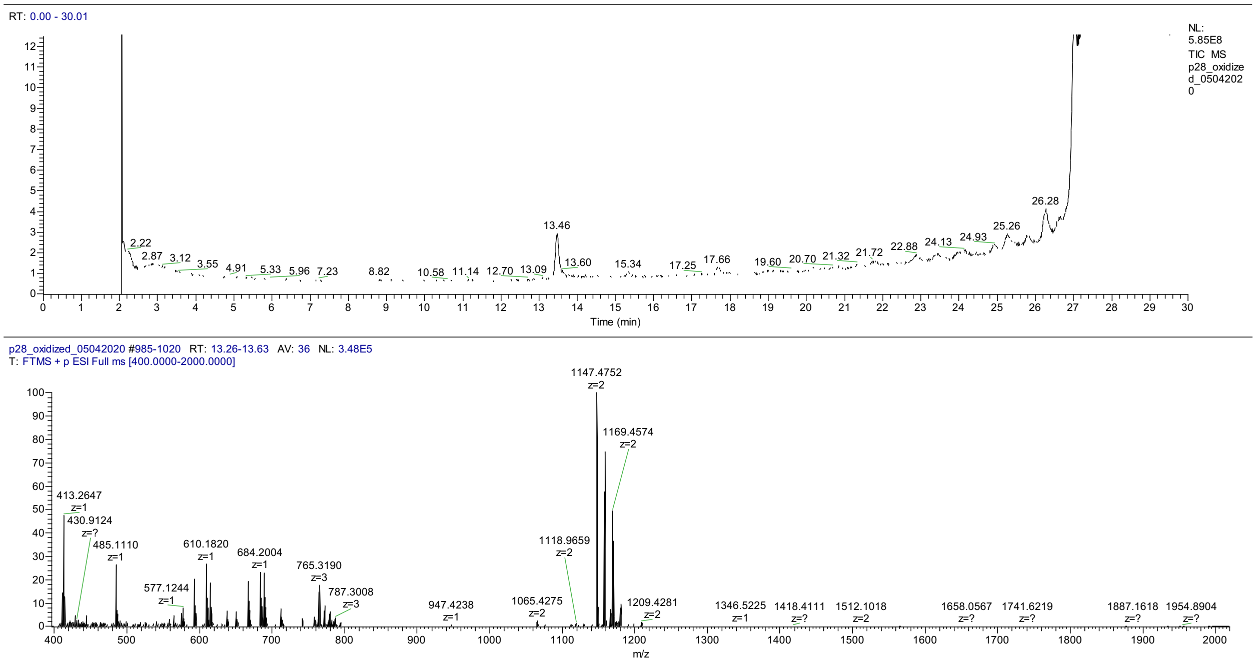


**Figure S8:** LC-MS data for P28. The TIC is shown on top and extracted data for the isolated peak on bottom.

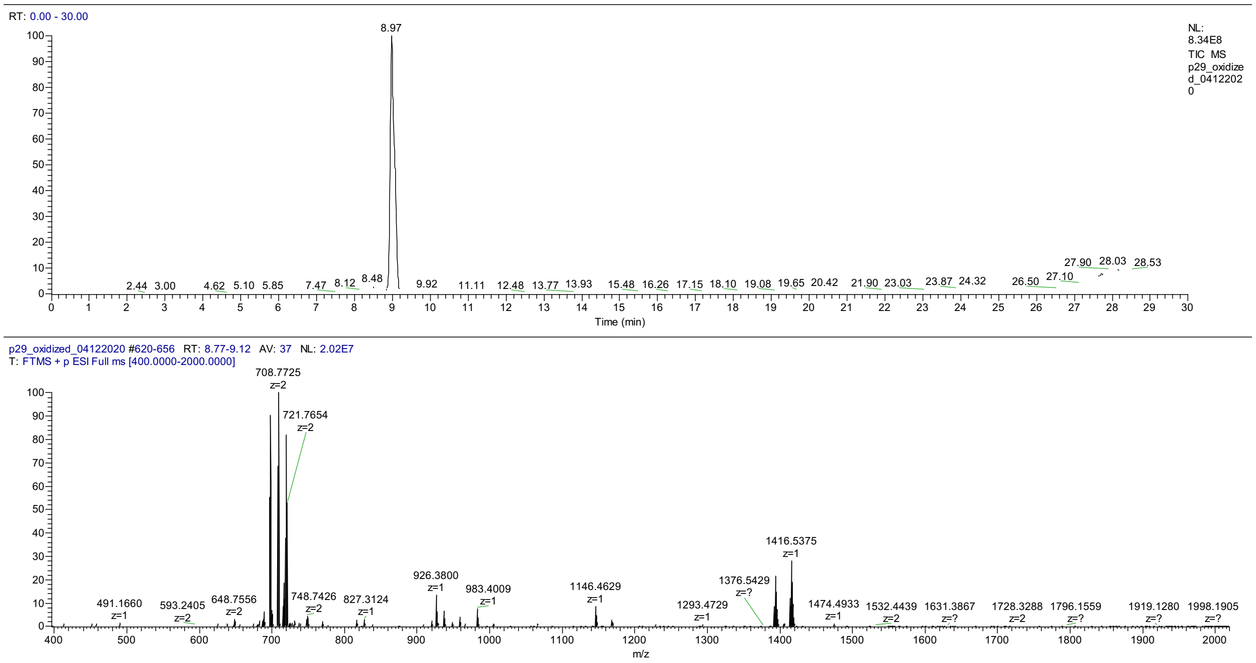


**Figure S9:** LC-MS data for P29. The TIC is shown on top and extracted data for the isolated peak on bottom.

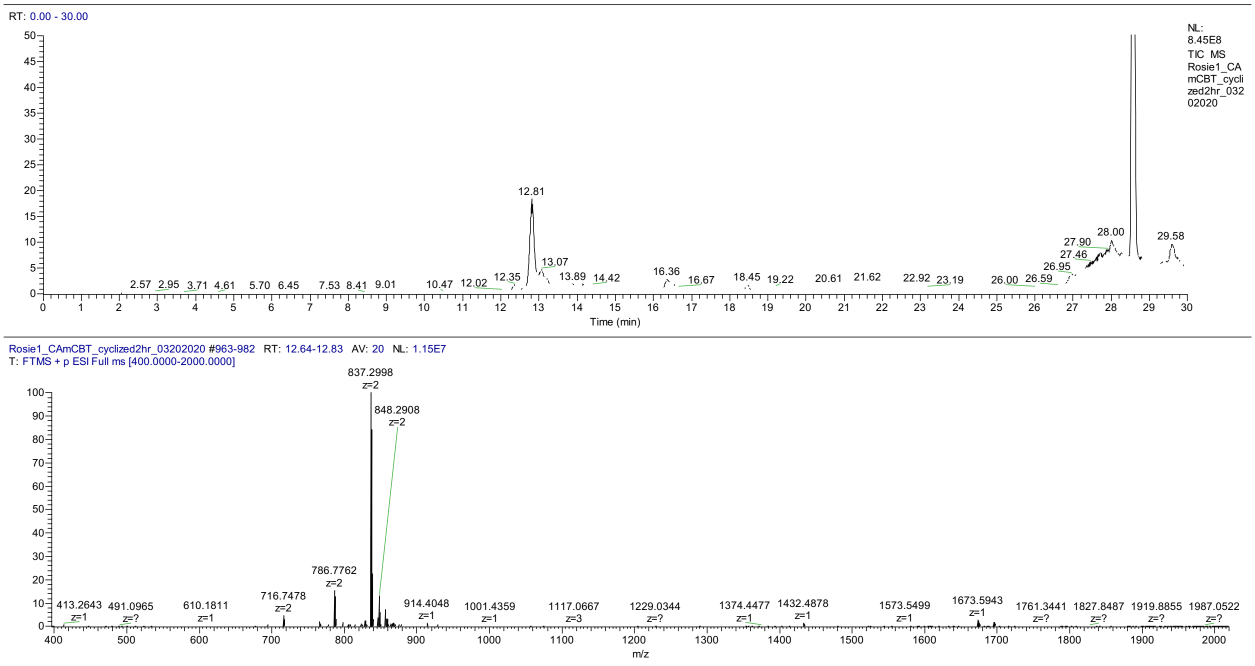


**Figure S10:** LC-MS data for P28S1. The TIC is shown on top and extracted data for the isolated peak on bottom.

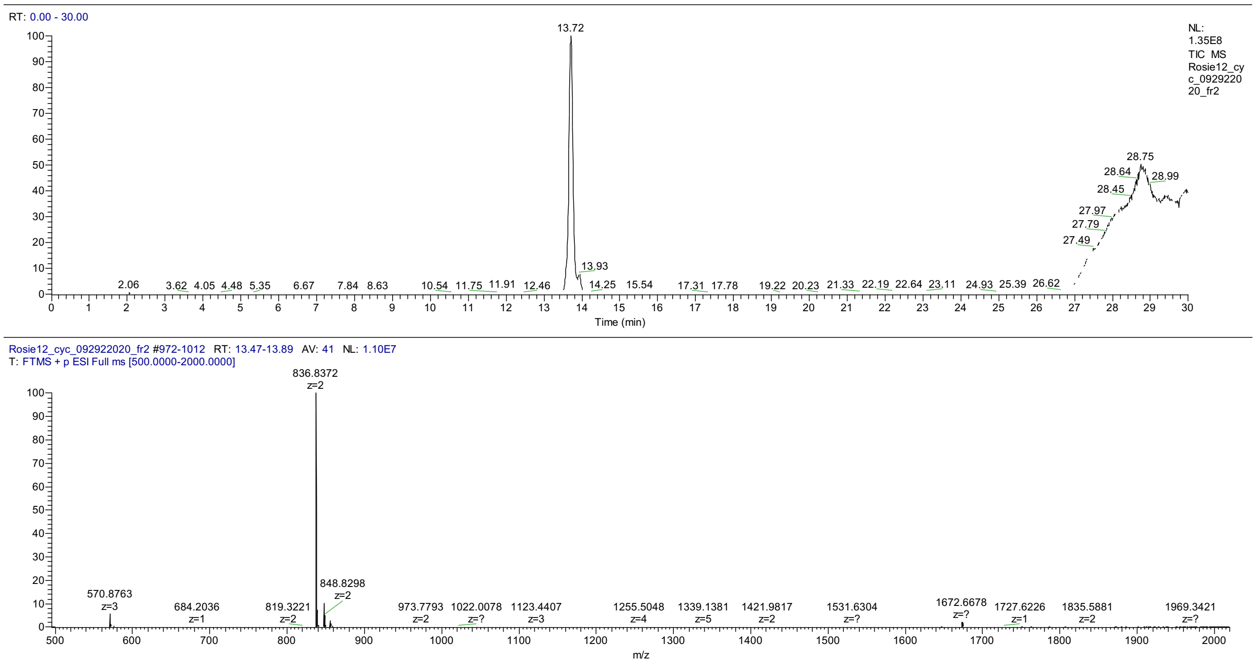


**Figure S11:** LC-MS data for P28S2. The TIC is shown on top and extracted data for the isolated peak on bottom.

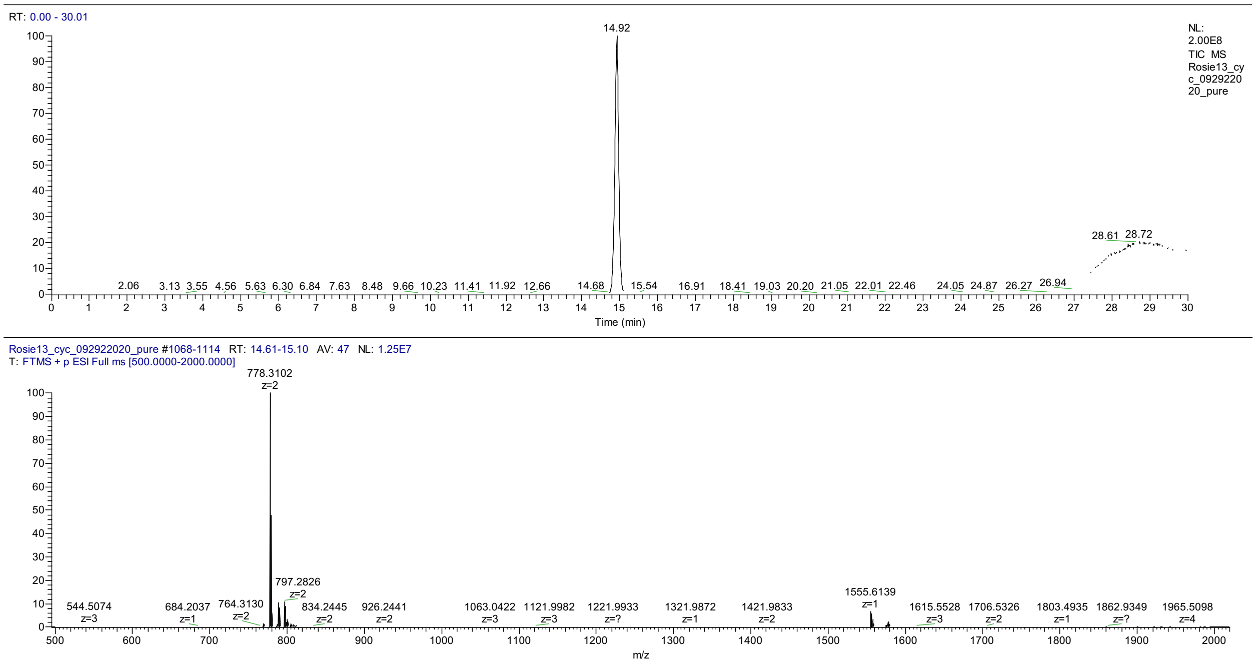


**Figure S12:** LC-MS data for P28S3. The TIC is shown on top and extracted data for the isolated peak on bottom.

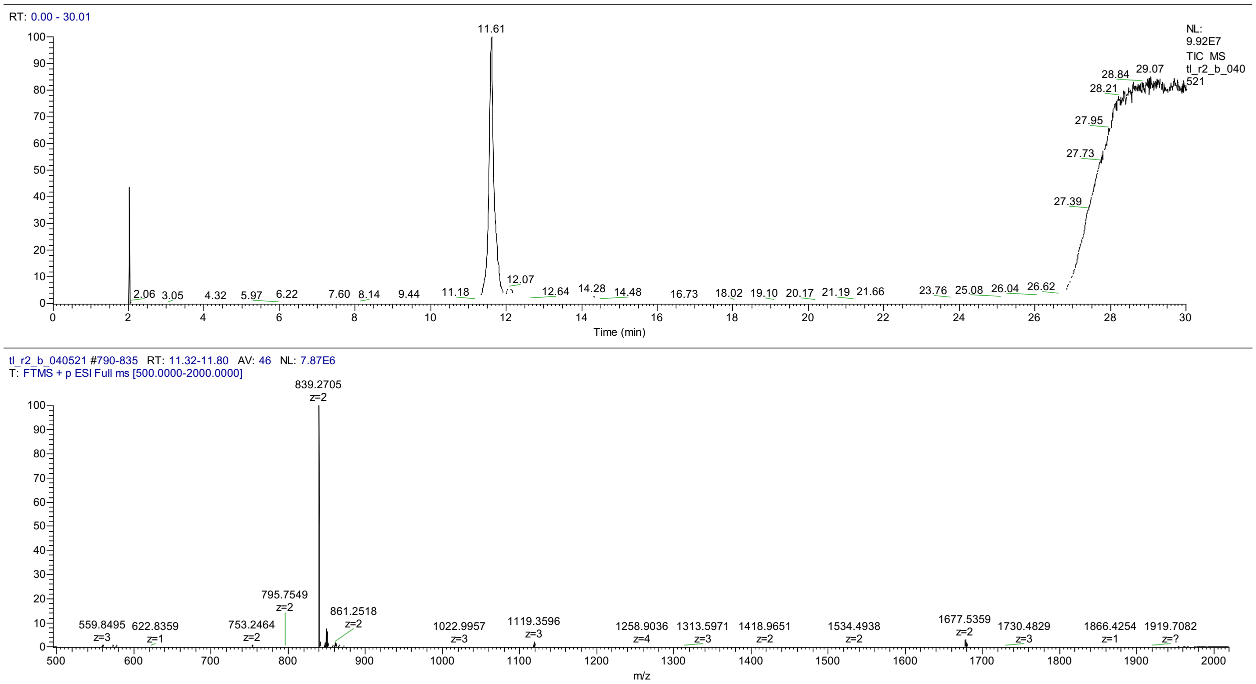


**Figure S13:** LC-MS data for P29S1. The TIC is shown on top and extracted data for the isolated peak on bottom.

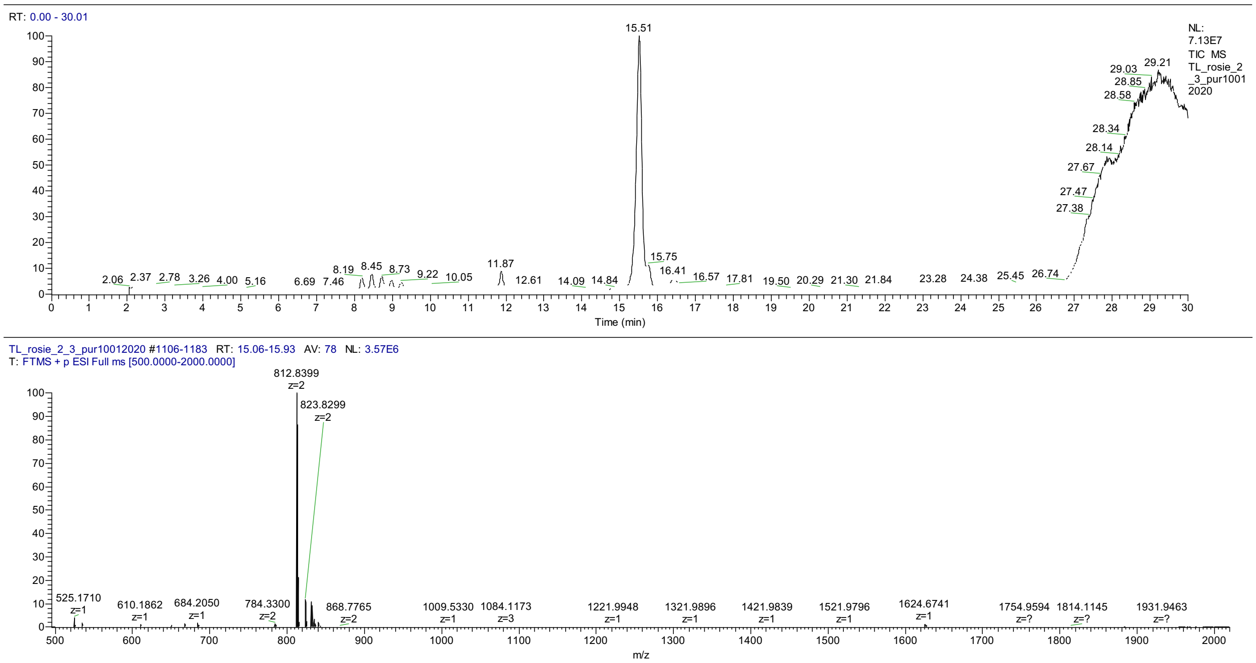


**Figure S14:** LC-MS data for P29S2. The TIC is shown on top and extracted data for the isolated peak on bottom.

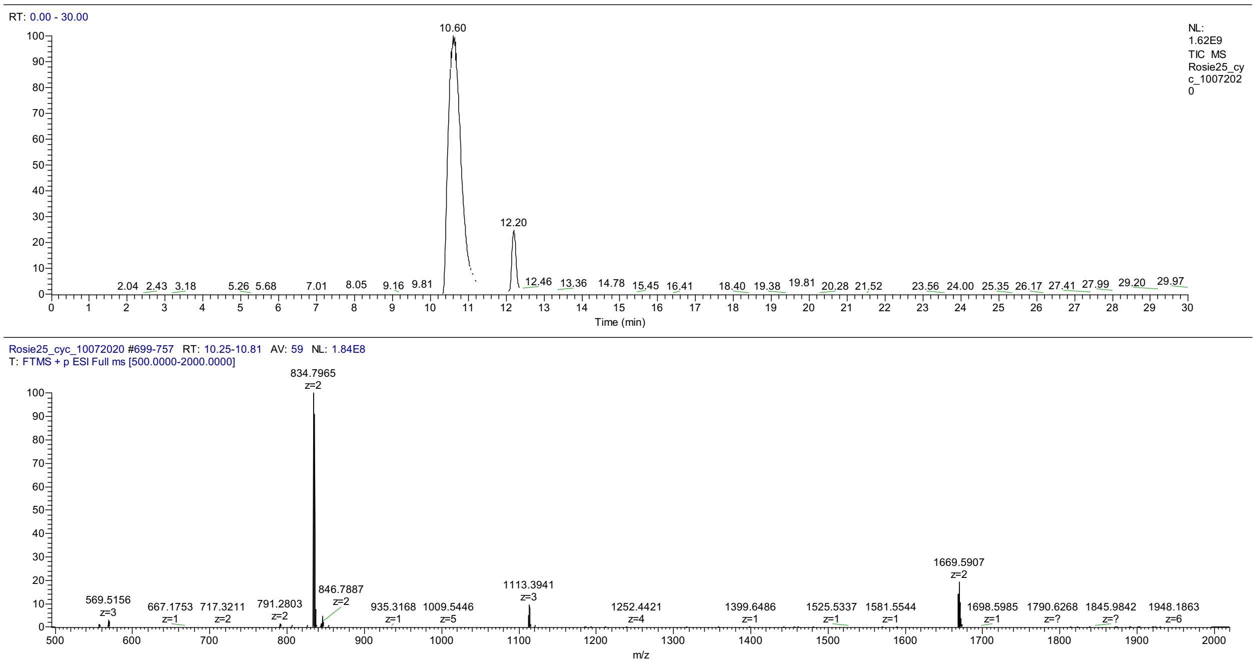


**Figure S15:** LC-MS data for P29S3. The TIC is shown on top and extracted data for the isolated peak on bottom.

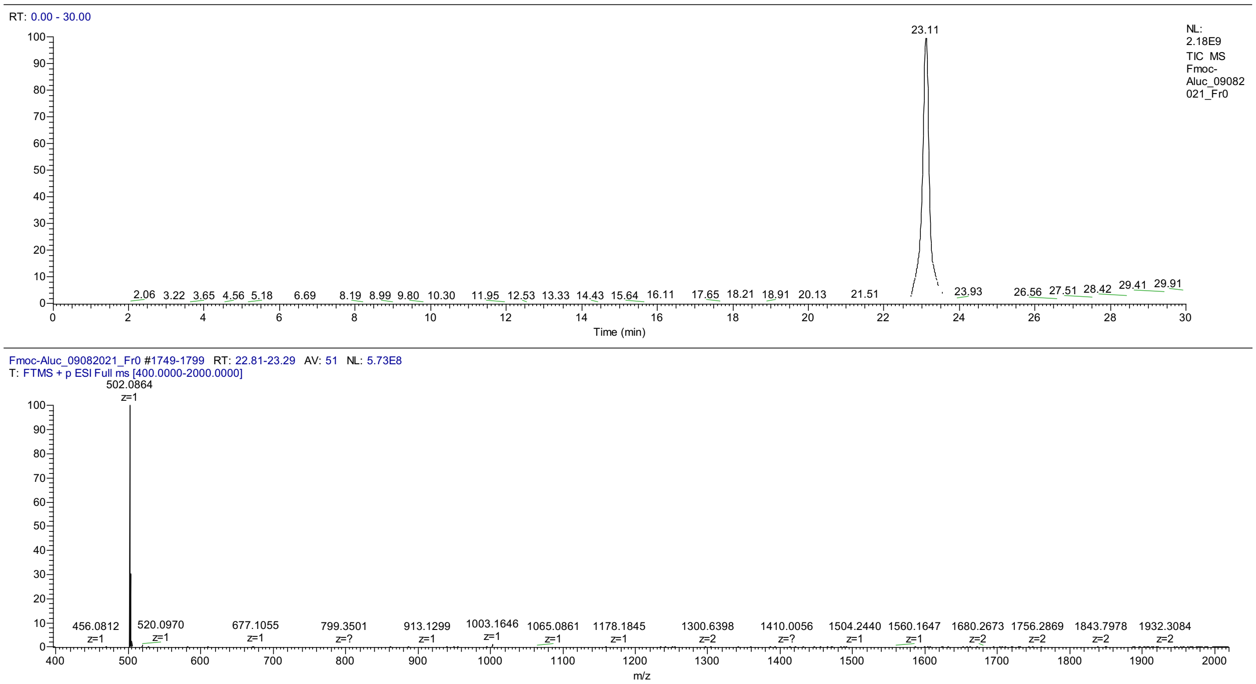


**Figure S16:** LC-MS data for Fmoc-Aluc. The TIC is shown on top and extracted data for the isolated peak on bottom.

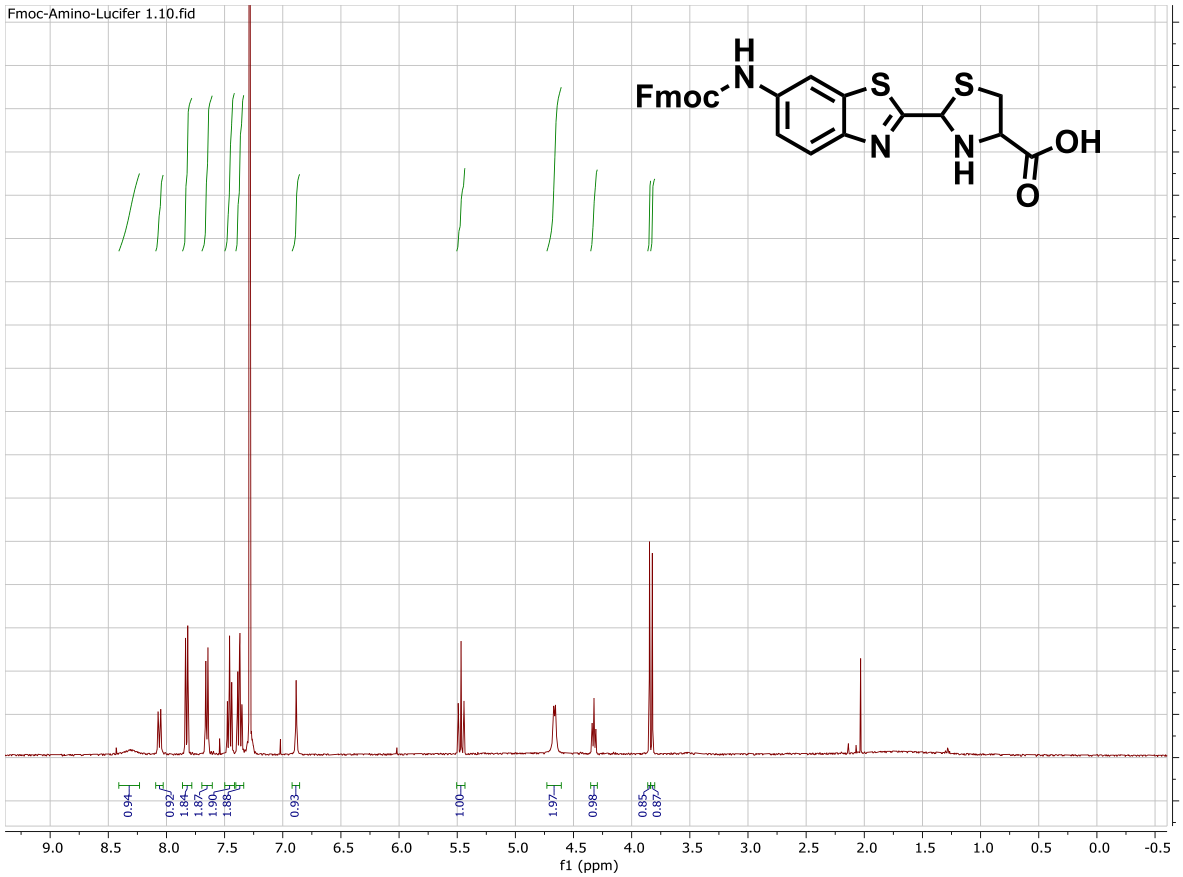


**Figure S17:** $^1$H NMR spectra for Fmoc-Aluc.

### SUPPLEMENTARY TABLES

**Table S1:** High-Resolution Mass Spectrometry Data for Synthesized Peptides

| Peptide | Expected Mass (Da) | Observed Mass (Da) |
|---|---|---|
| P28 | 2291.94 | 2291.94 |
| P29 | 1391.54 | 1391.55 |
| P28S1 | 1672.60 | 1672.59 |
| P28S2 | 1671.66 | 1671.67 |
| P28S3 | 1554.61 | 1554.61 |
| P29S1 | 1676.53 | 1676.54 |
| P29S2 | 1623.66 | 1623.68 |
| P29S3 | 1667.58 | 1667.59 |

**SUPPLEMENTARY SCRIPTS**

**Supplementary Script 1: Amino Acid Analysis**

```
library(microseq)
library(RColorBrewer)
library(dplyr)
library(stringr)
library(gplots)
NNK7Ffilt <-
 readFastq("/Users/traehampton/Documents/Research/Sequencing
 Results/Next Gen Sequencing/22005Wns_N22008/22005Wns_ZNRF3-
 R1_S1_L001_R1_001.fastq")
NNK7Rfilt <-
 readFastq("/Users/traehampton/Documents/Research/Sequencing
 Results/Next Gen Sequencing/21510Wns_N21170/21510Wns_12mer-
 negneg_S4_L001_R2_001.fastq")
#Define the following variables
libraryseq <- "GCCCAG.{54}GCGGCG.{6}" #change this regex to match
 specific library
beginning <- 19 #beginning of library in DNA string
lib <- 14 #number of codons in the library region
end <- beginning+lib*3
initialcodon <- beginning%/%3*4-1
endcodon <- initialcodon + lib*4
del <- beginning%/%3 #number of codons before library
aa <-
 c("A","C","D","E","F","G","H","I","K","L","M","N","P","Q","R","S","T",
 "V","W","Y","TAG")

#slices out matches that contain start followed by 24 bases to reverse
 primer
NNK7Ffilt21 <- gregexpr(libraryseq,NNK7Ffilt[,2],extract = TRUE)
NNK7Rrevcomp <- reverseComplement(NNK7Rfilt[,2],reverse = TRUE) #gives
 reverse complement of reverse reads
NNK7Rcompfilt21 <- gregexpr(libraryseq,NNK7Rrevcomp,extract = TRUE)

#this compares the forward and reverse strands, only allowing for one
 mismatch in the primers, no mismatches allowed in the library region
n <- length(NNK7Ffilt21)
NNK7Fgood <- vector()
for(i in c(1:n)){
  if(NNK7Ffilt21[[i]][1] == NNK7Rcompfilt21[[i]][1]){
    NNK7Fgood[i] <- NNK7Ffilt21[[i]][1]
```

```
  }
  else{
    split <- strsplit(c(NNK7Ffilt21[[i]][1],NNK7Rcompfilt21[[i]][1]),
 split = "")
    diff <- which(split[[1]] != split[[2]])
    if(length(diff) < 2 && length(diff) > 0){
      for(x in c(1:length(diff))){
        if(diff[[x]] < beginning || diff[[x]] > end){
          NNK7Fgood[i] <- NNK7Ffilt21[[i]][1]
        }
        else{
          NNK7Fgood[i] <- ""
        }
      }
    }
    else{
      NNK7Fgood[i] <- ""
    }
  }
}
NNK7Fgood <- as.data.frame(NNK7Fgood)
NNK7Fgood <- NNK7Fgood[!apply(is.na(NNK7Fgood) | NNK7Fgood == "", 1,
 all),]

#this separates nucleotides into codons
codons <- gsub("(...)", "\\1 \\2", NNK7Ffilt21)

#this creates dataframe of sequences with reads organized by frequency
seqcount <- as.data.frame(sort(table(codons), decreasing = TRUE))

#this generates a matrix that contains amino acids in library region
l <- length(codons)
AAs <- matrix(0,l,lib)
AA <- gregexpr("\\s(TT[TC])",codons,useBytes = FALSE)
  l <- length(AA)
  for(a in c(1:l)){
    l2 <- length(AA[[a]])
    for(b in c(1:l2)){
      value <- AA[[a]][b]
      if(value > initialcodon && value < endcodon){
        AAs[a,(value%/%4 - (del-1))] <- "F"
      }
    }
  }
```

```
AA <- gregexpr("(\\sTT[AG])|(\\sCT[GACT])",codons,useBytes = FALSE)
  l <- length(AA)
  for(a in c(1:l)){
    l2 <- length(AA[[a]])
    for(b in c(1:l2)){
      value <- AA[[a]][b]
      if(value > initialcodon && value < endcodon){
        AAs[a,(value%/%4 - (del-1))] <- "L"
      }
    }
  }
AA <- gregexpr("(\\sTC[GCAT])|(\\sAG[TC])",codons,useBytes = FALSE)
  l <- length(AA)
  for(a in c(1:l)){
    l2 <- length(AA[[a]])
    for(b in c(1:l2)){
      value <- AA[[a]][b]
      if(value > initialcodon && value < endcodon){
        AAs[a,(value%/%4 - (del-1))] <- "S"
      }
    }
  }
AA <- gregexpr("\\sTA[TC]",codons,useBytes = FALSE)
  l <- length(AA)
  for(a in c(1:l)){
    l2 <- length(AA[[a]])
    for(b in c(1:l2)){
      value <- AA[[a]][b]
      if(value > initialcodon && value < endcodon){
        AAs[a,(value%/%4 - (del-1))] <- "Y"
      }
    }
  }
AA <- gregexpr("\\sTAG",codons,useBytes = FALSE)
  l <- length(AA)
  for(a in c(1:l)){
    l2 <- length(AA[[a]])
    for(b in c(1:l2)){
      value <- AA[[a]][b]
      if(value > initialcodon && value < endcodon){
        AAs[a,(value%/%4 - (del-1))] <- "TAG"
      }
    }
  }
```

```
AA <- gregexpr("\\sTAA",codons,useBytes = FALSE)
  l <- length(AA)
  for(a in c(1:l)){
    l2 <- length(AA[[a]])
    for(b in c(1:l2)){
      value <- AA[[a]][b]
      if(value > initialcodon && value < endcodon){
        AAs[a,(value%/%4 - (del-1))] <- NA
      }
    }
  }
AA <- gregexpr("\\sTG[TC]",codons,useBytes = FALSE)
  l <- length(AA)
  for(a in c(1:l)){
    l2 <- length(AA[[a]])
    for(b in c(1:l2)){
      value <- AA[[a]][b]
      if(value > initialcodon && value < endcodon){
        AAs[a,(value%/%4 - (del-1))] <- "C"
      }
    }
  }
AA <- gregexpr("\\sTGA",codons,useBytes = FALSE)
  l <- length(AA)
  for(a in c(1:l)){
    l2 <- length(AA[[a]])
    for(b in c(1:l2)){
      value <- AA[[a]][b]
      if(value > initialcodon && value < endcodon){
        AAs[a,(value%/%4 - (del-1))] <- NA
      }
    }
  }
AA <- gregexpr("\\sTGG",codons,useBytes = FALSE)
  l <- length(AA)
  for(a in c(1:l)){
    l2 <- length(AA[[a]])
    for(b in c(1:l2)){
      value <- AA[[a]][b]
      if(value > initialcodon && value < endcodon){
        AAs[a,(value%/%4 - (del-1))] <- "W"
      }
    }
  }
```

```
AA <- gregexpr("\\sCC[GCAT]",codons,useBytes = FALSE)
  l <- length(AA)
  for(a in c(1:l)){
    l2 <- length(AA[[a]])
    for(b in c(1:l2)){
      value <- AA[[a]][b]
      if(value > initialcodon && value < endcodon){
        AAs[a,(value%/%4 - (del-1))] <- "P"
      }
    }
  }
AA <- gregexpr("\\sCA[CT]",codons,useBytes = FALSE)
  l <- length(AA)
  for(a in c(1:l)){
    l2 <- length(AA[[a]])
    for(b in c(1:l2)){
      value <- AA[[a]][b]
      if(value > initialcodon && value < endcodon){
        AAs[a,(value%/%4 - (del-1))] <- "H"
      }
    }
  }
AA <- gregexpr("\\sCA[AG]",codons,useBytes = FALSE)
  l <- length(AA)
  for(a in c(1:l)){
    l2 <- length(AA[[a]])
    for(b in c(1:l2)){
      value <- AA[[a]][b]
      if(value > initialcodon && value < endcodon){
        AAs[a,(value%/%4 - (del-1))] <- "Q"
      }
    }
  }
AA <- gregexpr("(\\sCG[GCAT])|(\\sAG[GA])",codons,useBytes = FALSE)
  l <- length(AA)
  for(a in c(1:l)){
    l2 <- length(AA[[a]])
    for(b in c(1:l2)){
      value <- AA[[a]][b]
      if(value > initialcodon && value < endcodon){
        AAs[a,(value%/%4 - (del-1))] <- "R"
      }
    }
  }
```

```
AA <- gregexpr("\\sAT[CAT]",codons,useBytes = FALSE)
  l <- length(AA)
  for(a in c(1:l)){
    l2 <- length(AA[[a]])
    for(b in c(1:l2)){
      value <- AA[[a]][b]
      if(value > initialcodon && value < endcodon){
        AAs[a,(value%/%4 - (del-1))] <- "I"
      }
    }
  }
AA <- gregexpr("\\sATG",codons,useBytes = FALSE)
  l <- length(AA)
  for(a in c(1:l)){
    l2 <- length(AA[[a]])
    for(b in c(1:l2)){
      value <- AA[[a]][b]
      if(value > initialcodon && value < endcodon){
        AAs[a,(value%/%4 - (del-1))] <- "M"
      }
    }
  }
AA <- gregexpr("\\sAC[GCAT]",codons,useBytes = FALSE)
  l <- length(AA)
  for(a in c(1:l)){
    l2 <- length(AA[[a]])
    for(b in c(1:l2)){
      value <- AA[[a]][b]
      if(value > initialcodon && value < endcodon){
        AAs[a,(value%/%4 - (del-1))] <- "T"
      }
    }
  }
AA <- gregexpr("\\sAA[CT]",codons,useBytes = FALSE)
  l <- length(AA)
  for(a in c(1:l)){
    l2 <- length(AA[[a]])
    for(b in c(1:l2)){
      value <- AA[[a]][b]
      if(value > initialcodon && value < endcodon){
        AAs[a,(value%/%4 - (del-1))] <- "N"
      }
    }
  }
```

```
AA <- gregexpr("\\sAA[AG]",codons,useBytes = FALSE)
  l <- length(AA)
  for(a in c(1:l)){
    l2 <- length(AA[[a]])
    for(b in c(1:l2)){
      value <- AA[[a]][b]
      if(value > initialcodon && value < endcodon){
        AAs[a,(value%/%4 - (del-1))] <- "K"
      }
    }
  }
AA <- gregexpr("\\sGT[GACT]",codons,useBytes = FALSE)
  l <- length(AA)
  for(a in c(1:l)){
    l2 <- length(AA[[a]])
    for(b in c(1:l2)){
      value <- AA[[a]][b]
      if(value > initialcodon && value < endcodon){
        AAs[a,(value%/%4 - (del-1))] <- "V"
      }
    }
  }
AA <- gregexpr("\\sGC[GACT]",codons,useBytes = FALSE)
  l <- length(AA)
  for(a in c(1:l)){
    l2 <- length(AA[[a]])
    for(b in c(1:l2)){
      value <- AA[[a]][b]
      if(value > initialcodon && value < endcodon){
        AAs[a,(value%/%4 - (del-1))] <- "A"
      }
    }
  }
AA <- gregexpr("\\sGA[TC]",codons,useBytes = FALSE)
  l <- length(AA)
  for(a in c(1:l)){
    l2 <- length(AA[[a]])
    for(b in c(1:l2)){
      value <- AA[[a]][b]
      if(value > initialcodon && value < endcodon){
        AAs[a,(value%/%4 - (del-1))] <- "D"
      }
    }
  }
```

```
AA <- gregexpr("\\sGA[AG]",codons,useBytes = FALSE)
  l <- length(AA)
  for(a in c(1:l)){
    l2 <- length(AA[[a]])
    for(b in c(1:l2)){
      value <- AA[[a]][b]
      if(value > initialcodon && value < endcodon){
        AAs[a,(value%/%4 - (del-1))] <- "E"
      }
    }
  }
AA <- gregexpr("\\sGG[GACT]",codons,useBytes = FALSE)
  l <- length(AA)
  for(a in c(1:l)){
    l2 <- length(AA[[a]])
    for(b in c(1:l2)){
      value <- AA[[a]][b]
      if(value > initialcodon && value < endcodon){
        AAs[a,(value%/%4 - (del-1))] <- "G"
      }
    }
  }
AAs <- as.data.frame(AAs)
#this gives unique amino acid sequences
UniqueAAs <- AAs %>% group_by_all() %>% count()
UniqueAAs <- UniqueAAs[order(-UniqueAAs$n),]
UniqueAAs <- UniqueAAs[apply(UniqueAAs,1,function(row) all(row !=
 0)),]
UniqueAAs <- na.omit(UniqueAAs)

#this counts sequences that have TAG codons, sequences that have more
 than one are only counted once
TAGreg <- regexpr("\\sTAG",codons)
TAGtable <- table(TAGreg)
percentTAG <-
 sum(TAGtable[2:length(TAGtable)])/length(codons)*100#percent of
 sequences containing TAG

#this creates a matrix of amino acid sequences that do not contain TAG
 codons
TAGpos <- which(AAs == "TAG")
TAGrow <- TAGpos%%nrow(AAs)
AAsnoTAG <- AAs[-TAGrow,]
```

```
#this creates heatmap for amino acid frequency per library position,
 change scale according to values
AAtable <- apply(AAs,2,function(x) table(factor(x,levels=aa)))
AAtable <- as.matrix(AAtable/length(codons))
colnames(AAtable) <- c(1:lib)
heatmapcolors <- colorRampPalette(brewer.pal(9,"Blues"))(100)
sc <- seq(0.0,0.6,by=0.006)
AAheatmap <- heatmap.2(AAtable, Rowv = NA, Colv = NA, col =
 heatmapcolors, density.info = "none", scale = "none", trace = "none",
 breaks = sc,  xlab = "Position in Library", ylab = "Codon", margins =
 c(3,4), dendrogram = "none")

#this creates projected heatmap based on NNK randomized codons
randomAAs <- matrix(0,21,lib,dimnames =
 list(rownames(AAtable),c(1:lib)))
randomAAs[c("A","G","P","T","V"),] <- 2/32
randomAAs[c("C","H","Q","N","K","Y","D","E","W","I","M","TAG","F"),]
 <- 1/32
randomAAs[c("L","S","R"),] <- 3/32
NNKheatmap <- heatmap.2(randomAAs, Rowv = NA, Colv = NA, col =
 heatmapcolors, density.info = "none", scale = "none", trace = "none",
 breaks = sc,  xlab = "Position in Library", ylab = "Codon", margins =
 c(3,4), dendrogram = "none")

#this creates heatmap showing bias from random, change scale with
 respect to range of values
lscale <- seq(-1,4,by=5/100)
librarybias <- (AAtable - randomAAs)/randomAAs
Biasheatmap <- heatmap.2(librarybias, Rowv = NA, Colv = NA, col =
 heatmapcolors, density.info = "none", scale = "none", trace = "none",
 breaks = lscale,  xlab = "Position in Library", ylab = "Codon",
 margins = c(3,4), dendrogram = "none")
librarybias <- as.data.frame(librarybias)

#this creates heatmap for AAsnoTAG
AAnoTAGtable <- apply(AAsnoTAG,2,function(x)
 table(factor(x,levels=aa)))
AAnoTAGtable <- as.matrix(AAnoTAGtable/nrow(AAsnoTAG))
colnames(AAnoTAGtable) <- c(1:lib)
heatmapcolors <- colorRampPalette(brewer.pal(9,"Blues"))(100)
sc <- seq(0.0,0.3,by=0.003)
AAheatmap <- heatmap.2(AAnoTAGtable, Rowv = NA, Colv = NA, col =
 heatmapcolors, density.info = "none", scale = "none", trace = "none",
```

```
breaks = sc,  xlab = "Position in Library", ylab = "Codon", margins =
 c(3,4), dendrogram = "none")

#writes csv files for uniqueAAs and bias heatmaps change path to make
 file
path <- "/Users/traehampton/Documents/Research/Sequencing Results/Next
 Gen Sequencing/21510Wns_N21170"
write.csv(UniqueAAs, paste(path,"/12mernegnegUniqueAAs.csv", sep =
 ""), row.names = F)
write.csv(TAGtable, paste(path,"/12mernegnegTAGtable.csv", sep = ""),
 row.names = T)
write.csv(librarybias, paste(path,"/12mernegnegLibraryBias.csv", sep =
""), row.names = T)
```

**Supplementary Script 2: Enrichment of Peptides**

```
#this gives overall enrichment of each peptide sequence by comparing
 Round 1 and Round 4 Library
library(prodlim)
path <- "/Users/traehampton/Documents/Research/Sequencing Results/Next
 Gen Sequencing/21510Wns_N21170"
lib <- 7
match <- row.match(UniqueAAsR1[,1:lib],UniqueAAsR3[,1:lib])
matchseq <- which(is.na(match)==FALSE)
percentenriched <-
 (UniqueAAsR3[match[matchseq],lib+1]/sum(UniqueAAsR3[,lib+1])-
 UniqueAAsR1[matchseq,lib+1]/sum(UniqueAAsR1$n))/(UniqueAAsR1[matchseq,
 lib+1]/sum(UniqueAAsR1$n))
enrichedseq <- UniqueAAsR3[match[matchseq],]
enrichedseq$enrichment <- percentenriched[,1]
enrichedseq <- enrichedseq[order(-enrichedseq$enrichment),]
enrichedseq <- as.data.frame(enrichedseq)
UniqueAAsR1$percent <- UniqueAAsR1$n/sum(UniqueAAsR1$n)*100
UniqueAAsR3$percent <- UniqueAAsR3$n/sum(UniqueAAsR3$n)*100
```
